# Testing alternative phylogenetic hypotheses for the tent tortoise species complex (Reptilia, Testudinidae) using multiple data types and methods

**DOI:** 10.1101/2020.11.02.364745

**Authors:** Zhongning Zhao, Neil Heideman, Jaco Oosthuizen, Margaretha D. Hofmeyr

## Abstract

We examined genetic differentiation in the highly polymorphic and taxonomically confusing tent tortoise (*Psammobates tentorius*) species complex in southern Africa, using three types of molecular markers (nDNA, mtDNA and microsatellite DNA) and morphological data. The Approximate Bayesian Computation based simulation analyses advocated an alternative phylogenetic hypothesis for the tent tortoise species complex, which was better and more inclusive in explaining its genealogical history. Based on the evidence derived from the sequence, microsatellite and morphology data, a four species scheme (among the seven mtDNA clades) appears to be the best taxonomic solution for the systematic puzzle of the *P. tentorius* species complex, namely, “C1+C4+C5+C7”, “C3”, “C2” and “C6”. The microsatellite datasets yielded similar genetic structure and gene flow patterns among the seven mtDNA clades in comparison to the sequence DNA. Evidence was found of possible hybridization between C1 and C2 in their intergradation zone, but not between C2 and C4. Results of the inbreeding analyses provided strong evidence of inbreeding in the eastern population of C1 and southern population of C2, which may be indicative of a bottleneck effect.

## INTRODUCTION

Africa is the centre of extraordinary chelonian diversity and high endemism, with 11 out of the 16 extant testudine genera found on this continent (Buhlmann et al., 2009; Mittermeier et al., 2015). Southern Africa harbours the world’s richest diversity of tortoise species, with a high level of endemism, 6 genera with 14 species occurring in this region (Hofmeyr et al., 2014; Mittermeier et al., 2015). Southern Africa is also regarded as a high priority region in terms of biodiversity conservation, since it possess 3 global biodiversity hotspots of both flora and fauna, namely, the Cape Floristic Region, Succulent Karoo and Albany thicket. Taxonomically the highly polymorphic tent tortoises (*Psammobates tentorius*, Bell, 1828) species complex of southern Africa is one of the world’s most confusing testudine groups (Branch, 2008; Hofmeyr et al., 2014; Rhodin et al., 2017), as manifested, amongst others, in the highly variable shape and colour patterns of its carapace and plastron. The widely distributed *P. tentorius* covers two of the 25 global biodiversity hotspots, namely, the Cape Floristic Region and Succulent Karoo.

Hewitt (1933 & 1934) described many species and subspecies based on regional character colour variations, Loveridge and Williams (1957) recognised only three subspecies, *Psammobates tentorius tentorius* (Bell, 1828), *Psammobates tentorius verroxii* (Smith, 1839) and *Psammobates tentorius trimeni* (Boulenger, 1886). This was also supported by Branch (2008), Hofmeyr et al., (2014). There are, however, still unresolved taxonomic complexities in some populations, particularly in *P. t. verroxii*, where the levels of polymorphism are the highest, warranting a re-evaluation of some of its synonymised taxa (Hofmeyr et al., 2014; Rhodin et al., 2017). There are substantial distribution range overlaps among the three subspecies, with seemingly hybrid individuals found in the intergradation zones, though many are believed to be misidentifications (Hofmeyr et al., 2014; Rhodin et al., 2017). Studying the *P. tentorius* species complex is also challenging as it generally occurs in low densities throughout its distribution range (Branch, 2008; Hofmeyr et al., 2014).

A recent phylogenetic study of the tent tortoise species complex by Zhao et al. (2020) based on six mtDNA loci and one nDNA marker revealed 7 distinct lineages (C1-C7). These findings placed clades C1, C4, C5 and C7 in the *P. t. tentorius* group, each occurring in a different geographic region generally isolated from the others by significant geographic barriers (see Fig. 1 and Supporting Information Figure S1-S2). The *P. t. verroxii* group was found not to be monophyletic; its populations north of the Orange River (C6) being genetically distinct from the populations south of the Orange River (C2). Furthermore, C2 and C6 were not sister taxa, implying that their designation as *P. t. verroxii* is not valid and should be reviewed (see Supporting Information Figure S1). The nDNA analysis, however, did not reveal a significant genetic difference between C2 and C6. These findings along with the uncertainty of the phylogenetic position of C6 (Zhao et al., 2020) therefore highlights the need for further investigation to resolve its taxonomic status, using additional markers. The phylogenetic position of a population from the Uniondale valley, not available to Zhao et al. (2020), also remains to be determined. Overall, the nDNA results showed some conflicts with the mtDNA findings regarding the placement of C3, and the relationship between C2 and C6. For illustrations showing the high phenotypic polymorphism in each clade, see Supporting Information Figures S1-S2.

**Figure 1.**
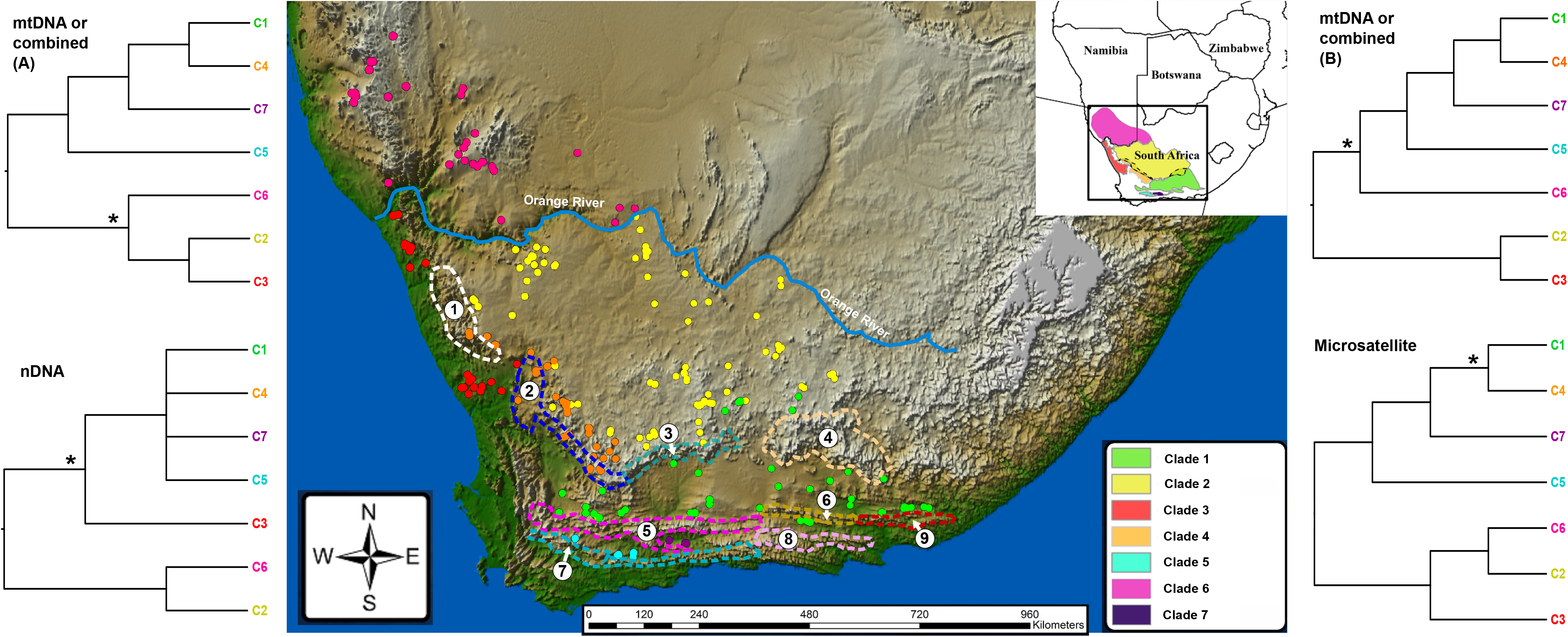
Summary of the four phylogenetic tree topologies generated with different analyses and datasets, namely, the two possible topologies retrieved from mtDNA and the combined sequence data (A and B), the nDNA-*PRLR* based tree, and a *Dsw* distance method using a neighbour-joining construction approach on the microsatellite dataset. “*” indicates weak support (BP < 70 or PP < 0.95). The map shows geographic distribution of the seven mtDNA clades throughout their range and samples used in the study are indicated in colours which correspond to their respective clades. The thin light blue belt represents a major river system between South Africa and Namibia. The major mountain barriers are indicated: 1) Kamiesberg, 2) Hantamberge and Roggeveldberge, 3) Nuweveldberge, 4) Sneeuberge and Kompasberg, 5) Swartberge and Rooiberg, 6) Grootrivierberge, 7) Langeberge, 8) Baviaanskloofberge, 9) Suurberge.

No investigation based on microsatellite and morphological markers has been conducted so far to scrutinize the above findings. In this study we attempted to provide further clarity on the matter by using multiple microsatellite DNA loci together with the sequence dataset of Zhao et al. (2020) and a comprehensive set of morphological data, in a finer scale genetic structure and morphological analysis of the species complex.

We used Approximate Bayesian Computation (ABC) to infer population history by carrying out simulations with our microsatellite DNA and mtDNA datasets under certain constrained scenarios (Beaumont et al., 2002). This was done in order to rank the likelihood of different scenarios and determine the most plausible cladogenic scenarios without being limited to the few scenarios of traditional phylogenetic analytical approaches. The traditional phylogenetic approach is usually restricted by limited scenario options and can be misleading in the interpretation of phylogenetic history, particularly when the study uses too few markers (Ballard & Rand, 2005; Cornuet et al., 2014). To address this issue and improve phylogenetic and genealogical reconstruction accuracy, we used ABC approach to determine the most likely genealogical scenario.

The influence of habitat on the morphology of organism has also been reported in other studies ((e.g., coastal fish by Farré et al. (2015); the snail, *Trochulus hispidus*, by Proćków et al. (2018)). Southern Africa is a region characterized by high levels of climatic, biome and vegetation heterogeneity (Mucina and Rutherford, 2011), which can be expected to drive the evolution of high levels of phenotypic plasticity in its fauna (Reed et al., 2011; Bonamour et al., 2019), as seems to have been the case in the *P. tentorius* species complex. Mismatches between the genotype and phenotype of species due to phenotypic plasticity under varying environmental conditions, are fairly common (Via and Lande, 1985). This is especially true in cryptic species complexes. Species richness can therefore be over- or under-estimated in studies based purely on morphological characters. A good example among tortoises would be the genus *Kinixys*. Six morpho-species previous recognised were shown to contain cryptic taxa, and that the actual number of *Kinixys* species was eight (Kindler et al., 2012). *Chersobius signatus* was initially believed to comprise two subspecies, *C. s. signatus* and *C. s. cafer*, based on morphology, but mtDNA and nDNA sequence data showed that the two “subspecies” were phylogenetically invalid (Daniels et al., 2010).

The aims of this study were, a) we infer population structuring of the *P. tentorius* species complex using both sequence and microsatellite data, and to determine the degree of congruence between the two types of markers, b) to carry out simulations with the combined DNA sequence and microsatellite datasets to determine the most plausible cladogenetic scenarios for the species complex, c) to determine the genetic diversity and inbreeding indices of clades, and potential gene flow among their populations from a conservation perspective, d) to investigate whether there was congruence between the findings based on molecular and morphological data, and e) to use the findings to make suggestions about the need for a taxonomic revision of the species complex.

## MATERIAL AND METHODS

### Sampling and DNA extraction

A total of 404 specimens of *P. tentorius* collected from 76 localities were used for the DNA sequencing and microsatellite genotyping. This comprehensively covered the species distribution range (Supporting Information Table S1). The collection and preservation of tissue and blood samples, and DNA extraction techniques, followed Zhao et al. (2020).

### mtDNA and nDNA amplification and sequencing

We used the Polymerase Chain Reaction (PCR) to amplify *12S* rRNA (*12S*), *16S* rRNA (*16S*), Cytochrome b (*Cyt-b*), NADH dehydrogenase subunit 4 (*ND4*), and the two *ND4* adjacent tRNA genes, *tRNA-His* and *tRNA-Ser*. For nDNA we used the fast-evolving Prolactin Receptor Coding gene (*PRLR*). The PCR set-up and reagents used followed Zhao et al. (2020). The optimal annealing temperatures and primers used for the sequence DNA markers are given in Supporting Information Table S2. All NCBI GenBank accession numbers of the sequence data are given in Supporting Information Table S1.

### Microsatellite DNA amplification, sequencing and genotyping

Extracted genomic DNA was used as template DNA in the PCR. Twenty-two microsatellite DNA loci, which were successful in previous testudines population genetics studies (Vamberger et al., 2001; Ciofi et al., 2002; Schwartz et al., 2003; Forlani et al., 2005; Paquette et al., 2005; Mandimbihasina et al., 2009; Orozco-terWengel et al., 2013), were used. They were tested on our target animal using PCR, with all forward primers labelled fluorescently with different dye probes (primer details and optimal annealing temperatures are given in Supporting Information Table S3). All PCR reactions were performed using KAPA2G Robust HotStart Readymix, USA, in a Bio-Rad T 100™ Thermal Cycler (Singapore) under the following parameters: an initial 5 min. denaturation step at 94 °C, followed by 35 cycles of 30 secs of denaturation at 94 °C, 30 secs of annealing (different optimal annealing temperatures were used depending on the loci, details given in Supporting Information Table S3), and a 1 min extension step at 72 °C, with a final 10 min extension step at 72 °C. The PCR annealing temperatures for the different microsatellite DNA loci were optimized using temperature gradient tests. Only loci showing a positive amplicon peak were used for further genotyping. All fragments were genotyped from chromatograms produced by an ABI 3500 genetic analyser.

We utilized multiplex PCR to enable the simultaneous amplification of multiple microsatellite DNA markers. We subdivided our microsatellite markers into six multiplex PCR groups according to the type of dye probes, fragment lengths and optimal PCR annealing temperatures, to maximize visualization of the genotyping and to minimize the interference among the different loci during genotyping (details given in Supporting Information Table S3). The PCR conditions for these six multiply mix reactions were the same as above for the single locus-based PCR. The annealing temperature for each multiple mix reaction is given in Supporting Information Table S3.

In order to verify the microsatellite repeat motif unit and genotype in the *P. tentorius* species complex, and to ensure that the amplicons obtained from the dye labelled primer pairs indeed came from the target region of interest, and not from off-target regions (caused by unspecific primer binding sites), we sequenced some individuals of the seven clades with the same primer pairs but without the dye labels (under the same PCR conditions applied during fragment analysis). The PCR products were electrophoresed in 1% agarose gel, visualized under UV light, and purified using a BioFlux PCR Purification Kit (Bioer Technology, China). Purified PCR products were cycle sequenced using BigDye (ABI PRISM® BigDye Terminator v3.1 Cycle Sequencing Kits, USA) and standard methods. The Big-Dye PCR products were purified with Zymo DNA Sequencing clean-up kits (Epigenetics Company, USA), prior to sequencing in an ABI 3500 genetic analyser.

We used PCR amplicon length for microsatellite genotyping and analyses throughout the study. In all cases, CONVERT (Glaubitz, 2004) and PGDSpider (Lischer and Excoffier, 2011) were used to convert datasets into different input file formats for different analyses. Sanger sequences were analysed using ABI Prism Sequencing Analysis software v.3.7 (Applied Biosystems), then aligned with MUSCLE v.3.2 (Edgar, 2004) and manually checked with MEGA v.7 (Kumar et al., 2016). Microsatellite DNA was genotyped with GeneMarker v.2.4 (Holland & Parson, 2011).

To ensure our microsatellite dataset did not contain null alleles, stuttering and large allele dropout (Oosterhout et al., 2004), we verified it using Micro-Checker (Oosterhout et al., 2004) to avoid misleading genotyping results. For all classes a combined probability of *p* < 0.05 was considered a significant indicator of the presence of null alleles.

### Population genetics analysis

#### Genetic structure

To determine genetic structure and significant clusters in the microsatellite dataset, STRUCTURE v 2.3 (Falush et al., 2003) was used to map the best cluster scheme by using Admixture as ancestry model to allow individuals the possibility of having mixed ancestry with the Bayesian algorithms. Analysis were run for 5 million generations using MCMC with the first 50 000 discarded as burn-in. All calculations were run for K = 1-11. STRUCTURE HARVESTER (Earl, 2012) was used to summarize outputs of STRUCTURE and to determine the best K value ((the one with the largest In Pr(X|K)). The program Clumpak (Kopelman et al., 2015) was used to visualize population structure across multiple cluster schemes (clustering under different K values).

Additionally, the Discriminant Analysis of Principal Components (DAPC, Jombart et al., 2010) was performed using the R package ‘adegenet’ (Jombart, 2008) of the program R v.3.5 (R Core Team, 2018). We first plotted a graph of the cumulative variance versus the number of Principle Components (PCs), initially retaining 400 PCs to initiate the analyses. We used the BIC criteria to determine the best k value (the number of clusters that should be retained), as visualized by a screen plot. We then used the a-score to measure the trade-off between discrimination power and over-fitting. The results of the a-score analysis were used to further optimize the number of PCs in the second DAPC analysis. Finally, we conducted a DAPC Cross-Validation test (using function “xvalDapc” to identify the ‘goldilocks point’ in the trade-off between retaining too few or too many PCs in the model. In order to test the genetic structure with finer scale, in this DAPC analysis, all individuals were divided into 11 populations (C1-central & western, C1-eastern, C2-northern, C2-southern, C3, C4, C5, C6-western, C6-eastern and the Uniondale population). Clade 1 was subdivided into a western and an eastern population. Clade 6 was also subdivided into a western and eastern population. Because both C1 and C6 showed clear subdivisions in Zhao et al. (2020). As for C2, the northern population was morphologically distinct from the southern population, we therefore subdivided it into a northern and southern population, although the mtDNA loci did not show them as clearly separate according to Zhao et al. (2020).

To infer relationships among lineages (C1-C7) with microsatellite markers, the POPTREE2 (Takezaki et al., 2009) was used to construct a distance-based unrooted neighbour-joining tree using microsatellite data. We selected *D*_*sw*_ distance methods (Shriver et al., 1995) with neighbour-joining tree construction methods using 1000 bootstraps to run the analyses.

SAMOVA v 2.0 (Dupanloup et al., 2002) was used to perform SAMOVA (Spatial Analysis of Molecular Variance) to identify populations that were geographically homogeneous and most likely differentiated from each other. It also assisted in identifying significant genetic barriers between these differentiated populations for both the mtDNA and microsatellite datasets. Different scenarios with K values ranging from 1 to 11 were tested independently, and the K scenario with the highest *F*_*CT*_ index was considered the best scenario. The SAMOVA incorporates a spatial dimension when considering the best genetic clustering scheme.

### Genetic diversity and gene flow

For both the mtDNA and microsatellite datasets, Arlequin v.3.5 (Excoffier & Lischer, 2009) was used to perform an Analyses of Molecular Variance (AMOVA) to evaluate population genetic structure of the seven mtDNA clades. The molecular diversity Theta (H) was calculated to measure polymorphism and allele diversity across the seven clades for the different microsatellite loci. The pairwise *F*_*ST*_ matrix, the average number of pairwise differences (π) within and between the seven clades, and Nei’s distance matrix (Nei and Li, 1979) were also calculated. The same indicators were estimated on the localities to evaluate the genetic diversity in regional scale (only localities with sample size n ≥ 3 were estimated). To evaluate the magnitude of gene flow and genetic divergence between the seven clades, we computed Slatkin’s linearized *F*_*ST*_ matrix (Slatkin, 1995) and relative divergence time between clades, based on the *F*_*ST*_ in Arlequin v 3.5 (Hudson et al., 1992).

We also used Migrate v 3.2 (Beerli, 2005 & 2009) to estimate the potential gene flow rate (M) and effective population size (Θ) with a Bayesian approach. The Migrate analysis of the microsatellite data assumed: i) constant population size through time with random fluctuation; ii) individuals mate randomly within each clade; iii) the mutation rate is constant through time and the same in all parts of the genealogy, and iv) migration rate is constant through time. We ran Migrate analyses to estimate the pairwise migration rates based on the seven mtDNA clades with one long MCMC chain, sampling every 1000 step for a total of 100 million genealogies after a burn-in of 100,000 steps in the chain. We used the Brownian motion stepwise mutation model. The Bayesian estimation of migration rate and population sizes was run with static heating (1, 1.5, 3, 10 000 000) and one long chain under a full model with all migration rates and population sizes. The M and Θ were both generated from *F*_*ST*_-calculations. The analysis results of Migrate were also used as prior estimations in later simulation analyses when testing alternative scenarios with the ABC approach.

### Hybridization and inbreeding

The R packages ‘poppr’ (Kamvar et al., 2014) and ‘adegenet’ were used to investigate possible hybridization among the seven mtDNA clades in potentially admixed individuals. This approach allows the automatic determination of the possible admixed individuals (possible hybridized individuals) using membership probabilities. We used the microsatellite dataset to perform this analysis. The threshold for detecting admixed individuals was set as less than 0.5 in predicted membership probability. The optimal number of PCs was determined in the same way as in the DAPC analysis.

In order to detect levels of inbreeding across different populations at fine scale, namely, C1 (western and central), C1 (eastern), C2 (northern), C2 (southern), C3, C4, C5, C6 (eastern), C6 (western) and C7, we performed an inbreeding detection analysis with microsatellite dataset using the R packages ‘ape’ (Paradis et al., 2004), ‘pegas’ (Paradis, 2010) and ‘adegenet’. Groups with frequency bars above 0.4 were considered as showing high inbreeding levels. We did not include the Uniondale samples in this analysis, since only two samples were available.

### Phylogenetic analysis and divergence time estimation

Because Zhao et al. (2020) did not conducted phylogenetic analysis based on combined dataset, to fill the gap, we combined the mtDNA dataset with the nDNA dataset to reconstruct the phylogeny. We performed a maximum likelihood analysis (ML) using RAxML v.8 (Stamatakis, 2014) with 1000 nonparametric bootstrap replications. The ML algorithm model selected was GTRCAT. We used the same partition scheme and parameter settings as Zhao et al. (2020), see Supporting Information Table S4. The substitution rate heterogeneity was allowed in different partitions. The accession numbers of outgroups sequences were given in (Supporting Information Table S5).

We also used BEAST v 2.4 (Bouckaert et al., 2014) with the StarBeast package (Drummond et al., 2012; Heled & Drummond, 2009) to carry out a Bayesian MSC phylogenetic analysis, generate a species tree and estimate the divergence time of each node. We used the same partition scheme, model test results and parameter settings as Zhao et al. (2020) (see Supporting Information Table S4). Regarding the prior setting for divergence time, seven nodes were constrained as monophyletic clades and calibrated with time intervals obtained from published studies (Cunningham 2002; Hofmeyr et al., 2017). A relaxed Log Normal clock was used during calculation. For each constrained node, we used a log-normal prior. Details of the constrained calibration nodes are given in Supporting Information Table S5). For the analysis we ran 50 million generations, sampling every 5000 generations and discarding the first 10% as burn-in. The results of the calibration dating analysis were used to estimate the divergence time priors (tau) in further ABC simulation analyses.

For the three new Uniondale museum samples, we were successful in obtaining 2 amplicons for the 12S gene but failed to gain any result from the other genes. We thus performed independent BI and ML analyses only on the 12S alignment including all samples from Zhao et al. (2020) and a few new sequences generated in this study. For the BI analysis we ran BEAST v2.4 for 50 million generations, sampling every 5000 generations and discarding the first 10% as burn-in. Parameter settings followed the model test results (see Supporting Information Table S4), as we assumed that the addition of the two new Uniondale samples was unlikely to influence the general parameter settings for the BI analysis. We used RAxML v.8 to perform the ML analysis with 100 nonparametric bootstrap replications under algorithm model GTRCAI.

### Bayesian simulations using the Approximate Bayesian Computations approach

In order to determine the most plausible genealogy scenario from the different tree topologies retrieved from different dataset, we used the Approximate Bayesian Computations (ABC) approach (Cornuet et al., 2008) implemented in DIYABC v 2.0 (Cornuet et al., 2014). This approach makes it possible to address much more complex and realistic biological situations using simulation methods. Comparing simulation results generated from different predefined scenarios using real datasets, enables detection of the most likely scenario from a wide range of scenarios, which cannot be tested with traditional likelihood based phylogenetic approaches. The ABC approach also allows the tracing of the past history of populations and species from different datasets. In this case, both sequence (mtDNA and nDNA) and microsatellite datasets were used to reconstruct the cladogenic history of the *P. tentorius* species complex to improve its accuracy.

To determine the most likely genealogical scenarios, we tested eight scenarios (see Fig. 2 for details), Scenario 1: assumed that C6 grouped with “C2+C3”, rather than “C1+C4+C7+C5”, and assumed “C1+C4+C7+C5” as the most ancestral lineage; this assumption was supported by the majority of the analyses using mtDNA and the combined datasets; Scenario 2: same as Scenario 1, except that “C6+C2+C3” was assumed as the most ancestral lineage; Scenario 3: assumed that C3 was sister to “C1+C4+C7+C5”, and C2 was sister to C6, “C3+C1+C4+C7+C5” was regarded as the most ancestral lineage. This scenario was supported by nDNA-*PRLR* and the morphological data; Scenario 4: the same as Scenario 3, except that “C6+C2” was assumed as the most ancestral lineage; Scenario 5: assumed that C3 was sister to “C2+C6”, rather than group with “C1+C4+C7+C5”, “C1+C4+C7+C5” was the most ancestral lineage. This scenario was supported by the microsatellite data; Scenario 6: same as Scenario 5, but assumed “C6+C2+C3” as the most ancestral lineage; Scenario 7: as indicated in Zhao et al. (2020), for the mtDNA, and the combined data in this study, the uncertainty of the placement of C6 sometimes resulted in C6 grouping with “C1+C4+C7+C5” rather than “C2+C3”. This scenario assumed “C6+C1+C4+C7+C5” as the most ancestral lineage; and Scenario 8: same as Scenario 7, except that it assumed “C2+C3” as the most ancestral lineage.

**Figure 2.**
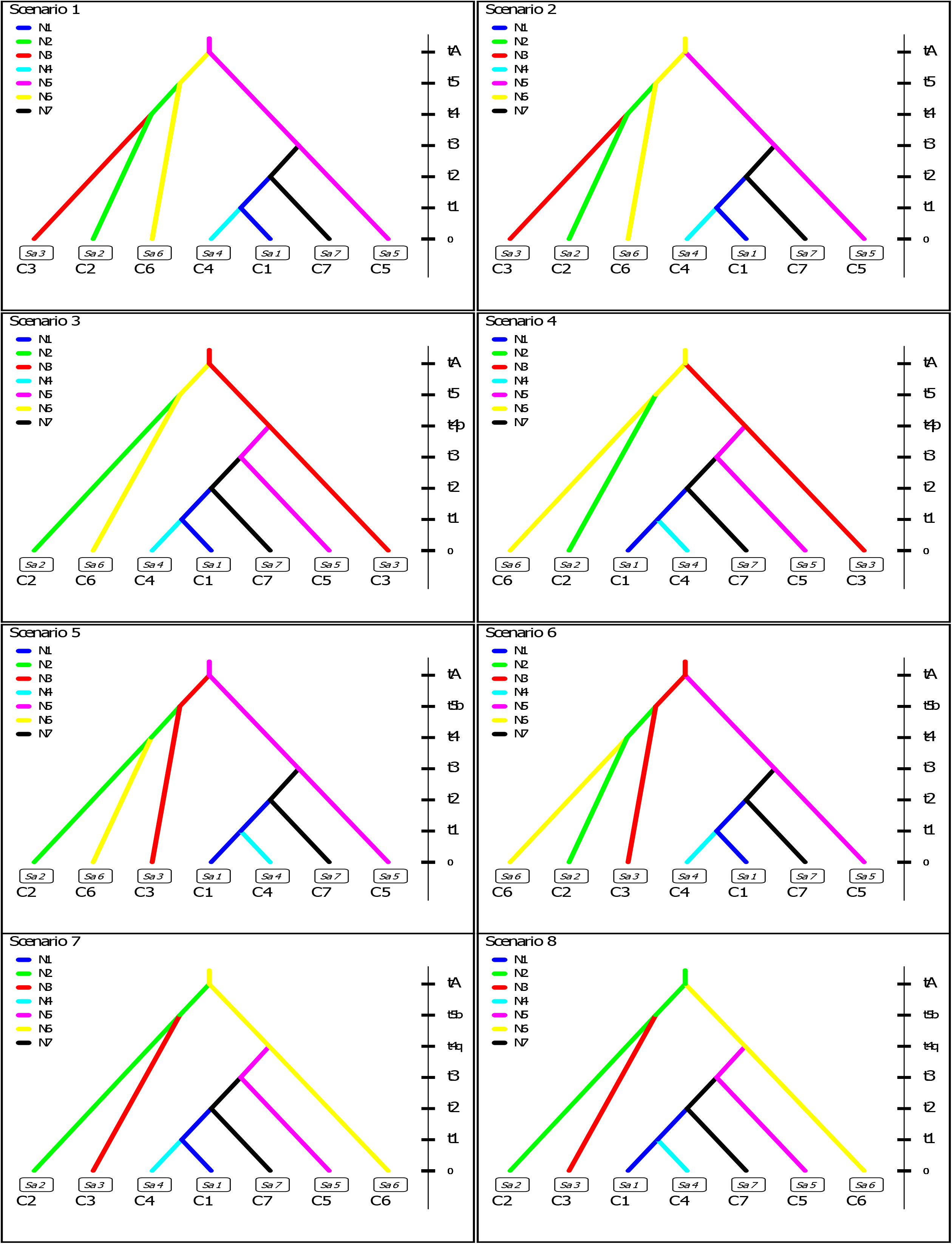
The eight possible scenarios used in the Approximate Bayesian Computation (ABC) analyses. Each of these scenarios was constructed based on different possible scenarios retrieved from different phylogenetic analyses with the different types of molecular markers (nDNA, mtDNA and microsatellite DNA). Each pre-defined genealogical assumption was used in simulations run in the ABC analysis. In each pre-defined scenario, the clade names for corresponding branches are given below them.

To define a proper prior for improving run accuracy, the effective population sizes (*Ne*) of the seven clades were estimated from the microsatellite dataset results generated by Migrate v.3.2. The divergence time priors (tau) were estimated from the four possible topologies, the mtDNA tree, the nDNA tree, the combined tree and the microsatellite-based tree topology for different scenarios. The prior settings for tau were, tA > t5 > t4 > t3 > t2 > t1 > 0, t4p > t3, t5b < tA and t4q > t3 (see Fig. 2), following the StarBeast dating results.

We divided the dataset into three groups, the mtDNA dataset, the nDNA dataset and the microsatellite dataset, each with independent mutation models. The priors for the mtDNA dataset group were specified as, mean mutation = 7.00 × 10^⎕3^ substitution/site/million years, with range 1.0 × 10^−3^ - 1.0 × 10^−2^ substitution/site/million years. Whilst, for the nDNA dataset group, the mutation rate priors were specified as, mean mutation = 3.26 × 10^⎕4^ substitution/site/million years, with range 1.0 × 10^−4^ - 1.0 × 10^−3^ substitution/site/million years, based on the evolutionary rate estimated for turtles by Lourenco et al. (2013). The mutation model used was HKY+I+G with invariant sites = 50% and shape of gamma = 1. We used the suggested default settings for the prior setting scheme in the microsatellite group. In the sequence dataset group (mtDNA and nDNA), we chose the number of haplotypes, the number of segregating sites, the mean pairwise differences and Tajima’s D for the one sample summary statistics, and the mean pairwise differences (W) and *F*_*ST*_ as objects for the two sample summary statistics. For the microsatellite dataset group, we selected mean number of alleles, mean genetic diversity and mean size of variance for the one sample summary statistics, and mean genetic diversity and *F*_*ST*_, classification index, shared allele distance and (dμ)_2_ distance for the two sample summary statistics. We simulated 9.0 x 10^6^ datasets in the initial run following the system suggested for the optimal number of runs for obtaining both computationally and statistically robust results, before further processing using the ABC analysis.

For the final summary analysis of the simulation results, all 8 million simulated datasets in the reference table were used. We first performed a pre-evaluative prior-scenario combination analysis to evaluate whether our observed data fell into the simulated prior range of each scenario, using a preliminary Principle Component Analysis (PCA). If the observed dataset fell into the simulated prior scenario range, it meant that the simulated results were reliable, and that sampling was adequate and even. We then computed the posterior probability of each scenario in order to determine the most likely scenario, using linear discriminant analysis. The results were plotted to visualize and rank the posterior scores of the different simulated scenarios. To visualize and quantify the posterior distribution of the different simulation parameters, as well as to ensure prior settings were adequate, we estimated the posterior distribution of each parameter and plotted it independently. Finally, we performed a model checking analysis on each simulated scenario to ensure that our observed dataset fell into the range of priors and posteriors of the simulated results of each PCA scenario.

### Morphological analyses

Detailed information regarding the specimen and data collection, grouping variables and data partitions, and characters used in the morphometric (continuous) and phenotypic (discrete) datasets are given in the Supplementary Materials (Supporting Information Tables S6-S7, Figures S3-S4). Prior to carrying out any of the analyses, the datasets were tested for normality using the Shapiro-Wilks test in the program SPSS v.20. If the requirements for normality were not met, the variables were log-transformed and again checked for normality.

### Sexual dimorphism

Prior to performing multivariate analyses with both morphometric and phenotypic datasets to determine whether the seven clades were morphologically different, we checked for sexual dimorphism, to avoid gender bias influencing our multivariate analyses. Sexual size dimorphism was determined using two-way PERMANOVA with 10000 permutations on the morphometric and phenotypic dataset, respectively, with PAST (Paleontological Statistics) v.3.08 (Hammer et al. 2001).

### Distinguishing morphologically among the clades

To remove the effect of body size, each log-transformed variable was regressed against log CLS (straight carapace length) as covariate using a general linear model in SPSS v.20. The studentized linearized residuals were retrieved and used as input data in further multivariate analyses.

To investigate how the clades clustered relative to each other, a Discriminant Function Analysis (with the Generalized Discriminant Analysis model) and a Principal Component Analysis (with correlation matrix) were performed on the linearized residuals in STATISTICA v.8. Only the scores of the first two discriminant functions or principal components were retrieved for analyses and exported to SPSS v.20 or PAST v.3.08 to generate better quality scatterplots for visualization. Since significant sexual dimorphism was found, the continuous datasets were partitioned by sexes prior to the DFA, and separate analyses performed.

The one-way PERMANOVA analysis with 10000 permutations in PAST was used to determine whether the seven clades were significantly different from each other in terms of their morphology (morphometric and phenotypic data, separately). Because of significant sexual dimorphism, separate analyses were carried out for each sex.

## RESULTS

### Microsatellite DNA amplification, sequencing and genotyping

The coverage of the mtDNA sequencing was generally good (see Supporting Information Table S1), however, we failed to amplify nDNA-*PRLR* for the majority of carcass samples. The microsatellite DNA genotyping results showed that three of the 21 loci of the *P. tentorius* species complex failed to amplify during PCR, namely, GP19, GP30 and GAL75, while the rest of the loci produced amplification results (Supporting Information Table S8). The loci RAD932, Maucas22, TEST10, GAL100 and GAL127 showed comparatively large numbers of failed amplifications in certain individuals. The Micro-Checker analysis results revealed significant null allele signals for loci RAD932, Maucas22 and Test10 (*p* < 0.001 in all cases). The analyses failed for the GAL100 and GAL127 loci, as both had too many missing alleles. We therefore excluded them from the dataset before the final analysis commenced. We therefore used the 14 loci dataset (Supporting Information Table S9) to carry out further analyses.

### Population genetics analysis

#### Genetic structure

In the DAPC analysis, the BIC criterion suggested four clusters (K=4) as optimal clustering scheme (Supporting Information Figure S5). The a-score optimisation retrieved fourteen PCs as optimal number for further DAPC analysis (Supporting Information Figure S5). The cross-validation test confirmed that the number of PCs retrieved for the analyses was sufficient (Supporting Information Figure S5). The DAPC scatterplot (Fig. 3A) revealed four candidate species, species A consisted of C1 (western and eastern populations), C4, C5 and C7; species B comprised C3; species C included the eastern and western populations of C6; species D contained the northern and southern populations of C2.

**Figure 3.**
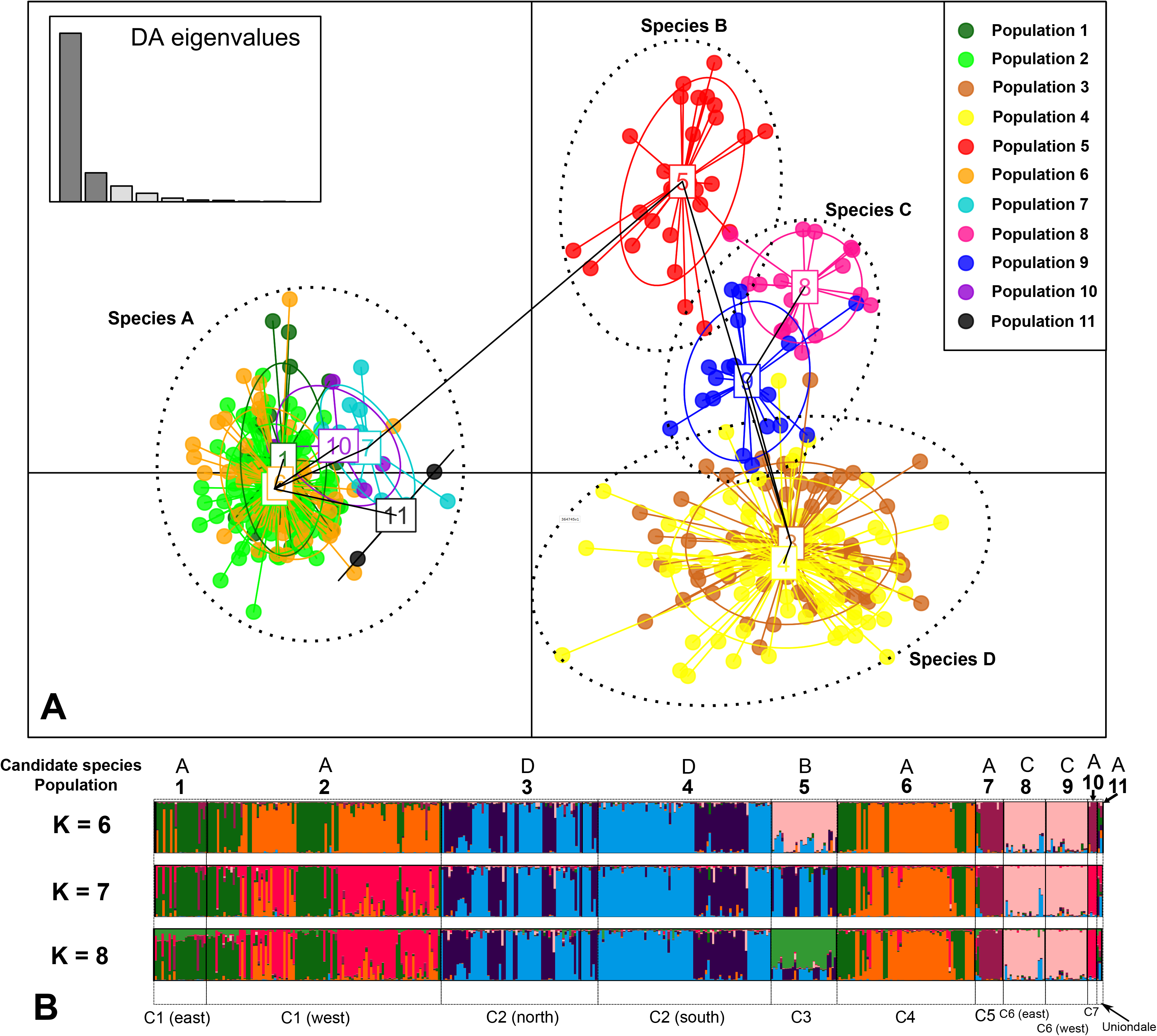
**A**: Scatterplot of the DAPC analysis among 11 populations, using 14 microsatellite DNA loci, with the “four candidate species” scheme also given. Population 1: C1 (eastern), Population 2: C1 (western), Population 3: C2 (northern), Population 4: C2 (southern), Population 5: C3, Population 6: C4, Population 7: C5, Population 8: C6 (eastern), Population 9: C6 (western), Population 10: C7 and Population 11: Uniondale population. **B**: The putative clustering schemes of the 11 populations generated from STRUCTURE analyses (K = 6 – 8) with corresponding “four candidate species” scheme.

The STRUCTURE HARVESTER results revealed K = 6 as the most likely number of clusters. We nevertheless also present the results for K = 7 and K = 8, to compare how the different clusters match the seven mtDNA clades and the 11 populations (Fig. 3B). The STRUCTURE results did not exhibit a huge incongruence with the seven mtDNA clades, showing five STRUCTURE clusters that corresponded to the four candidate species retrieved from the DAPC analysis. The difference between C3 and C6 was, however, not clear under K = 6. The differences between C3 and C6 were, however, significant under runs K = 8 to K = 11. Furthermore, the STRUCTURE results failed to distinguish between C1 and C4, and C5 and C7. None of the subdivisions at within clade level ((C1 (eastern), C1 (western), C2 (northern), C2 (southern), C6 (eastern) and C6 (western)) revealed significant differences.

### Spatial genetic structure

The SAMOVA generally suggested 6 clusters as the optimal spatial-scale genetic structure for the microsatellite DNA dataset, since the over-split event was at its minimum level in each clade, though the *F*_*CT*_ value was not the best at K = 6 (Supporting Information Table S10). In the optimal clustering scheme (see Fig. 4), the eastern but not the western population of C6 differed significantly from C2, while C1 and C4 did not differ from each other. The spatial clusters C3, C5 and C7 perfectly matched their mtDNA clades.

**Figure 4.**
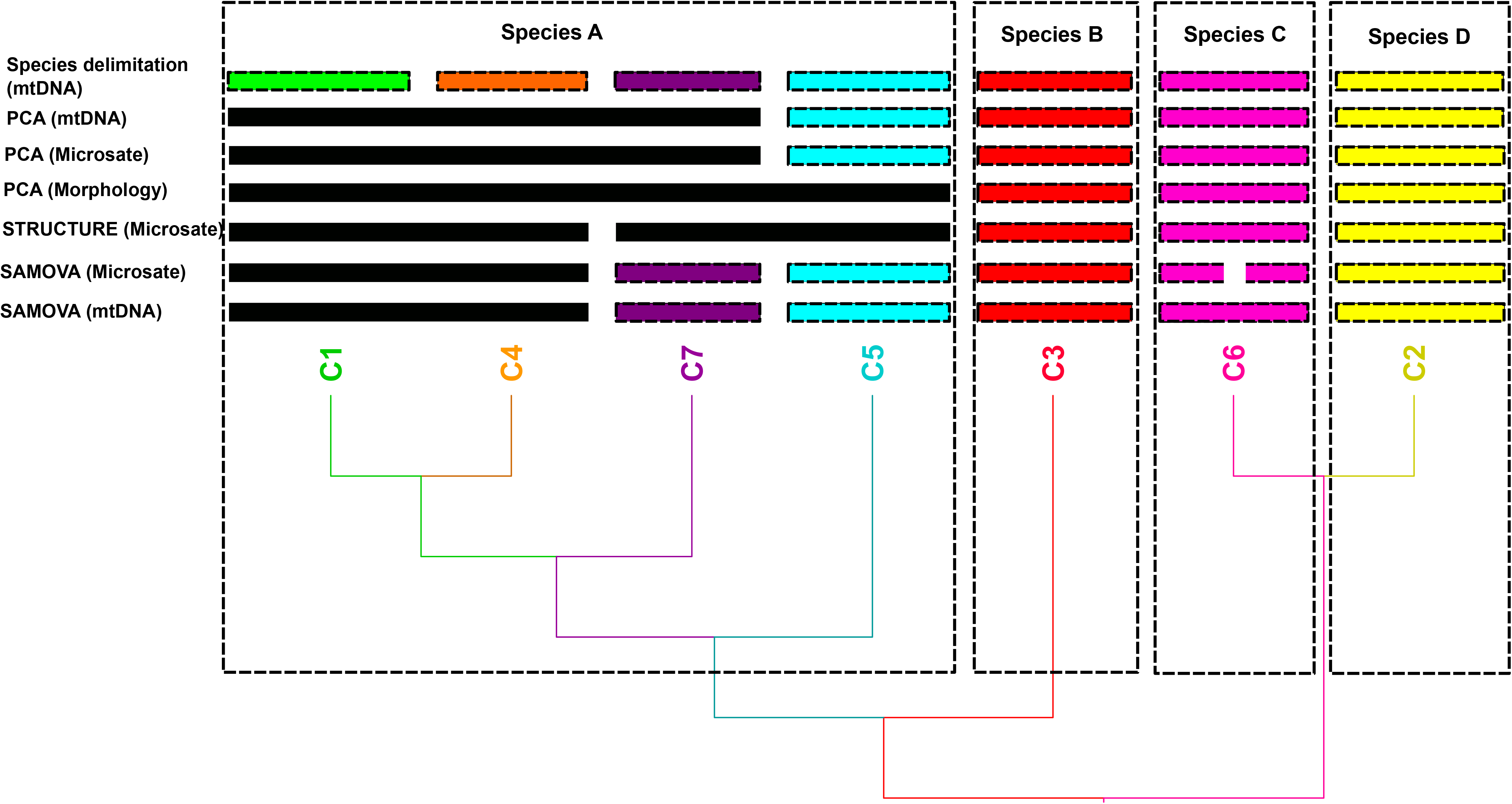
The bar plot shows the optimal “four species” scheme under summary of different delimitation approaches, together with the most likely genealogy (the most likely topology advocated by the ABC simulation analysis based on sequence and microsatellite data). The colour scheme is the same as the mtDNA clades indicated elsewhere.

For the mtDNA dataset, the SAMOVA also retrieved 6 spatial clusters (K=6), but the cluster allocations differed from that of the microsatellite DNA dataset (Fig. 4). No significant difference was found between C1 and C4, while the rest of the clades were significantly different as distinct spatial clusters.

### Genetic diversity and gene flow potential

The AMOVA results of the microsatellite DNA dataset revealed that 12.38% of the total genetic variation was from among the seven clades (Va: df=6, sum of squares=453.42 and variance components=0.69), 23.95% of the total variation came from within each clade (Vb: df=395, sum of squares=2458.02, and variance components=1.33) and 63.67% of the variation came from within individuals (Vc: df=402, sum of squares=1427.5, and variance components=5.58). The fixation indices *F*_*IS*_, *F*_*ST*_ and *F*_*IT*_ were 0.27, 0.13 and 0.36, respectively. The standard genetic diversity of microsatellite DNA was highest in C1 and C4, and lowest in C7 (Table 1). The genetic diversity index, Theta (H), also revealed the highest genetic diversity in C1 and C4, and the lowest in C7 (Table 1). The same trend was found in the allele diversity. The matrix of the average number of pairwise differences suggested a major divergence between the two major groups, “C1+C4+C5+C7” (southern) and “C2+C3+C6” (northern). In southern group, the lowest pairwise difference was found between C1 and C4, and the highest between C5 and C7, whilst, in northern group, the highest difference was found between C3 and C6, and the lowest between C2 and C6. The highest within-group pairwise distance was found in C1 and C4. The Uniondale population was closest to C1 (Fig. 5). The Slatkins linearized *F*_*ST*_ matrix, tau matrix and *F*_*ST*_ matrix generally showed identical trends to the pairwise distance matrix. At population level (Table 1), the eastern population of C1 exhibited much lower genetic and allele diversity than the western population. The genetic diversity levels of the northern and southern populations of C2 did not show a great difference, the southern population having slightly higher genetic diversity (Theta, H) than the northern population. The genetic diversity between western and eastern populations of C6 was almost identical.

**Figure 5.**
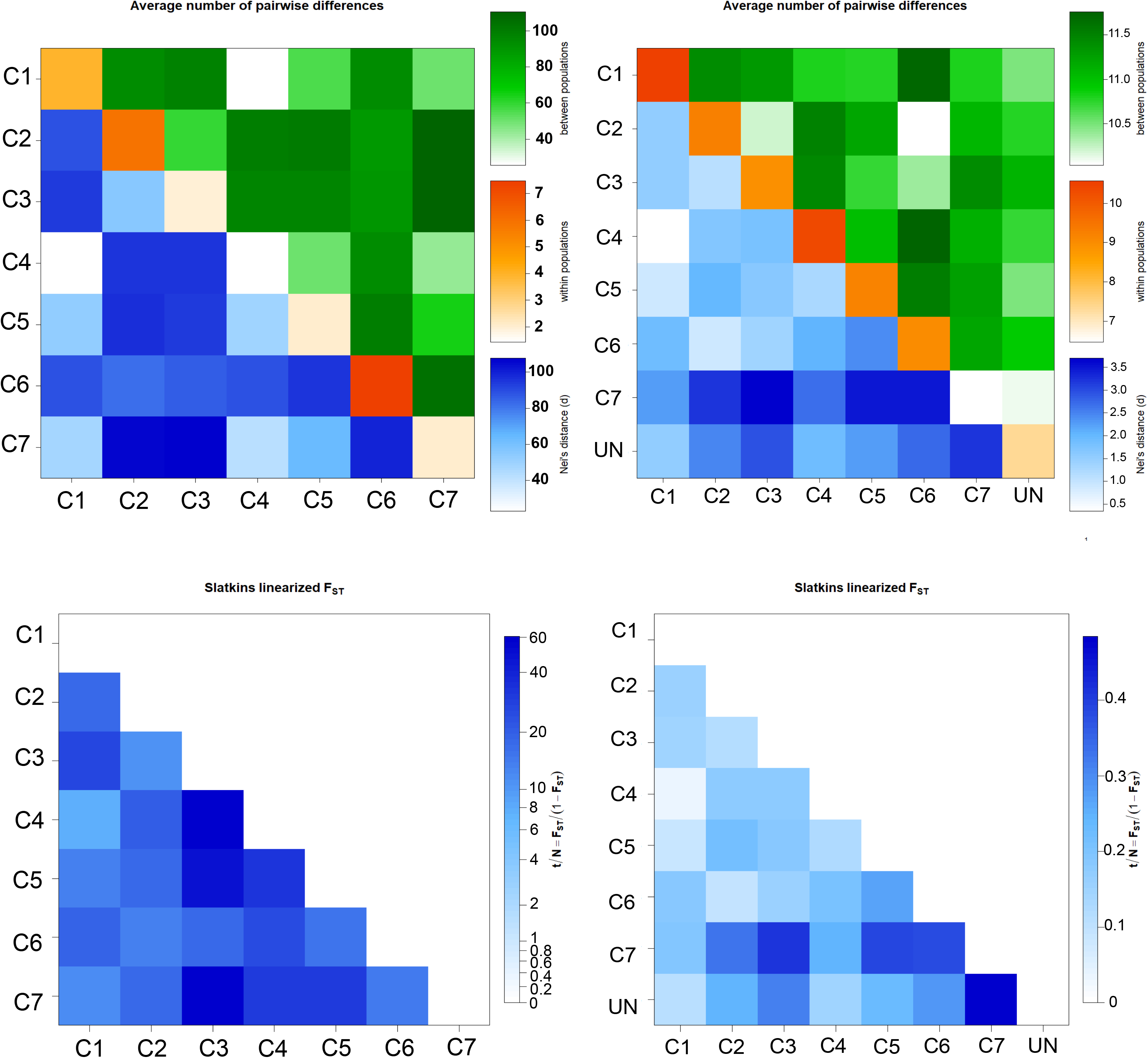
Top left: the average number of pairwise differences among and within different clades derived from the Arlequin analyses using the mtDNA dataset. Top right: the average number of pairwise differences among and within different clades derived from the Arlequin analyses using the microsatellite DNA dataset. Bottom left: the gene flow indicator-Slatkin’s linearized *F*_*ST*_ matrix among different clades derived from the Arlequin analyses using the mtDNA dataset. Bottom right: the gene flow indicator-Slatkin’s linearized *F*_*ST*_ matrix among different clades derived from the Arlequin analyses using the microsatellite DNA dataset. “UN” refers to the Uniondale population.

**Table 1.** AMOVA analysis results showing sample size (N), polymorphic sites, number of haplotypes (no. Haplotype), gene diversity, genetic diversity (Theta, H) and number of alleles (no. of alleles) among different populations and clades, using the mtDNA and microsatellite DNA datasets.

| mtDNA |  |  |  |  |  |
| --- | --- | --- | --- | --- | --- |
| population/clade | N | polymorphic sites | No. Haplotype | gene diversity | Theta (H) |
| C1 | 123 | 39 | 52 | 0.95 | 17.34 |
| C1_western | 104 | 28 | 38 | 0.93 | 13.73 |
| C1_eastern | 19 | 11 | 14 | 0.71 | 1.78 |
| C2 | 151 | 52 | 64 | 0.97 | 32.72 |
| C2_northern | 79 | 25 | 22 | 0.91 | 10.2 |
| C2_southern | 72 | 30 | 33 | 0.95 | 18.13 |
| C3 | 32 | 13 | 11 | 0.68 | 1.64 |
| C4 | 63 | 9 | 12 | 0.59 | 1.06 |
| C5 | 12 | 10 | 7 | 0.83 | 4.04 |
| C6 | 38 | 38 | 22 | 0.95 | 16.92 |
| C6_western | 19 | 19 | 9 | 0.87 | 7.9 |
| C6_eastern | 19 | 19 | 12 | 0.9 | 15.9 |
| C7 | 4 | 4 | 3 | 0.83 | 4.02 |
| overall | 423 | 288 | 171 | 0.98 | 49.38 |

| Microsatellite DNA |  |  |  |  |  |  |
| --- | --- | --- | --- | --- | --- | --- |
| population/clade | N | polymorphic loci | No. Haplotype | gene diversity | Theta (H) | No. allele |
| C1 | 122 | 14 | 244 | 0.75 | 3.22 | 26 |
| C1_western | 100 | 14 | 200 | 0.76 | 3.21 | 25 |
| C1_eastern | 22 | 13 | 44 | 0.58 | 2.46 | 10 |
| C2 | 141 | 14 | 282 | 0.66 | 2.05 | 22 |
| C2_northern | 60 | 12 | 120 | 0.61 | 2.02 | 15 |
| C2_southern | 81 | 14 | 162 | 0.67 | 2 | 17 |
| C3 | 28 | 14 | 56 | 0.64 | 1.77 | 9 |
| C4 | 59 | 13 | 118 | 0.75 | 3.17 | 17 |
| C5 | 12 | 12 | 24 | 0.65 | 1.99 | 6 |
| C6 | 36 | 13 | 72 | 0.64 | 1.8 | 13 |
| C6_western | 18 | 13 | 36 | 0.62 | 1.65 | 9 |
| C6_eastern | 18 | 12 | 36 | 0.61 | 1.56 | 9 |
| C7 | 4 | 11 | 8 | 0.51 | 0.92 | 3 |
| mean | 402 | 14 | 804 | n/a | 2.04 | 12 |

The Arlequin genetic analysis results of the mtDNA dataset generally found the highest genetic diversity in C1, C2 and C6, and the lowest in C3 and C4 (Table 1). The mtDNA dataset also detected a major divergence between southern and northern groups across the different matrices. At population level, the eastern population of C1 showed much lower genetic diversity than the western population. In C2 the genetic diversity was higher in the southern than northern population and in C6 the eastern population exhibited significantly greater genetic diversity than the western population. At locality level, the intergradation zones (Calvinia, Loeriesfontein, Richmond and Victoria West) where two clades coexist exhibited substantially higher allele and genetic diversity than expected. Besides these, localities Prince Albert, Matjiesfontein and Steytlerville also exhibited higher genetic diversity than the rest (see Table 1 and Supporting Information Figure S6 and Table S11).

In the microsatellite DNA dataset, the Migrate analysis results suggested that the effective population size (Theta, H) was similar across the seven clades (Supporting Information Table S12). We therefore did not differentiate between the prior settings for the different Theta values among the clades in the ABC simulation analysis. The posterior distribution plots for the different migration parameters were normally distributed, and no significant “double peak” was observed. In terms of gene flow, the highest rate was found between C1 and C2, and C1 and C4, whilst the lowest was found between C1 and C5, C1 and C7, and C5 and C7. The gene flow rate between C2 and C4, and between C2 and C6 was also relatively high, but comparatively low between C3 and C6. These intergradation zones also revealed higher genetic diversity in terms of average number of alleles, polymorphic loci and gene diversity. Lainsburg, Prince Albert, Matjiesfontein and Loxton also exhibited relatively higher genetic diversity level than the rest of the localities. Recent migration events were discovered by Migrate analysis between C1 and C2, C1 and C4, C3 and C4, C4 and C5, and C4 and C7, but not between C2 and C6, and C3 and C6.

### Hybridization and inbreeding

The DAPC admixture analysis detected 10 suspected hybridized individuals (Supporting Information Figure S7), seven of which came from the Victoria West and Richmond area, suggesting hybridizations between C1 and C2. The remaining three were assigned as crossovers among C4, C5 and C7. There was no any admixture signal detected as expected in the intergradation zone between C2 and C4.

The eastern population of C1 and southern population of C2 had a significantly higher inbreeding signal than the rest populations, with the majority of samples falling into a group with an inbreeding coefficient > 0.4 (Supporting Information Figure S8).

### Phylogeny and divergence time estimation

In the single locus 12S based phylogeny, both ML and BI approaches, however, suggested C1 as sister group to the Uniondale samples (Supporting Information Figure S9), with strong support (BP > 70, PP > 0.95). The ML results for the combined dataset (mtDNA+nDNA) generated a tree topology similar to the mtDNA based phylogenetic tree of Zhao et al. (2020), except for some differences in branch lengths (Supporting Information Figure S10). The different nodes were generally well supported, (BP > 70), except the uncertainty of the placement of C6, again.

The microsatellite DNA distance-based neighbour-joining tree (Fig. 1) showed a similar tree topology to the mtDNA and combined (mtDNA+nDNA) tree, except that the relationship between C2, C3 and C6 was different. Clade 2 and C6 formed a sister group, and the relationship between C3 and (C2+C6) was supported. The topology of (C1+C4)+C7)+C5 was nonetheless well supported in the microsatellite based phylogeny, which was identical to the mtDNA and combined (mtDNA+nDNA) trees. Each node had strong support (BP > 70), except the one between C1 and C4.

The BI calibration dating results based on the combined dataset revealed similar divergence time estimations between the gene tree and species tree (Supporting Information Figure S11). The calibrated species tree generally suggested that the radiation of the *P. tentorius* species complex dated back to the Mid-Miocene (about 14.17 Mya). The divergence time between C2+C3 and C6 was given as 12.18 Mya, and between C2 and C3, as 8.23 Mya. The branch-off time between C5 and (C1+C4)+C7 was dated as 7.67 Mya, which was similar in age to the node between C2 and C3, but slightly more recent. Clade 7 split from C1+C4 about 5.49 Mya, while C4 diverged from C1 about 3.97 Mya. The dating results of outgroup nodes exhibited high congruence with the calibration dating results of Hofmeyr et al. (2017). These dating results were used in prior setting for tau in ABC analysis.

### Bayesian simulations of alternative scenarios

The results of the ABC simulation analyses advocated scenario 4 as the most likely one for explaining the real genealogical history of the *P. tentorius* species complex (see Fig. 6 and Supporting Information Figure S12), rather than the expected scenario 1 or 2, which were well supported by the previous derived mtDNA and mtDNA+nDNA based phylogeny. The ABC simulation results were closer to the tree topology retrieved from the nDNA dataset and microsatellite DNA loci. This suggested C6 as the most likely ancestral lineage and advocated that C2 and C6 were sister groups and clustered as a major branch (the *P. t. verroxi* complex). Clade 3 (*P. t. trimeni*) clustered with all the *P. t. tentorius* lineages (C1+C4)+C5)+C7, forming a second major branch. The PCA results of priors and posteriors (Supporting Information Figures S13-S14) revealed that the actually observed dataset fell completely into the range of simulated datasets for all scenarios, which implies adequate sampling with a high level of confidence in its reliability. When comparing the parameter plots between priors and posteriors, we generally found good matches between their distributions (Supporting Information Figure S15). It therefore showed that the sampling during simulations was adequate and the simulation results reliable.

**Figure 6.**
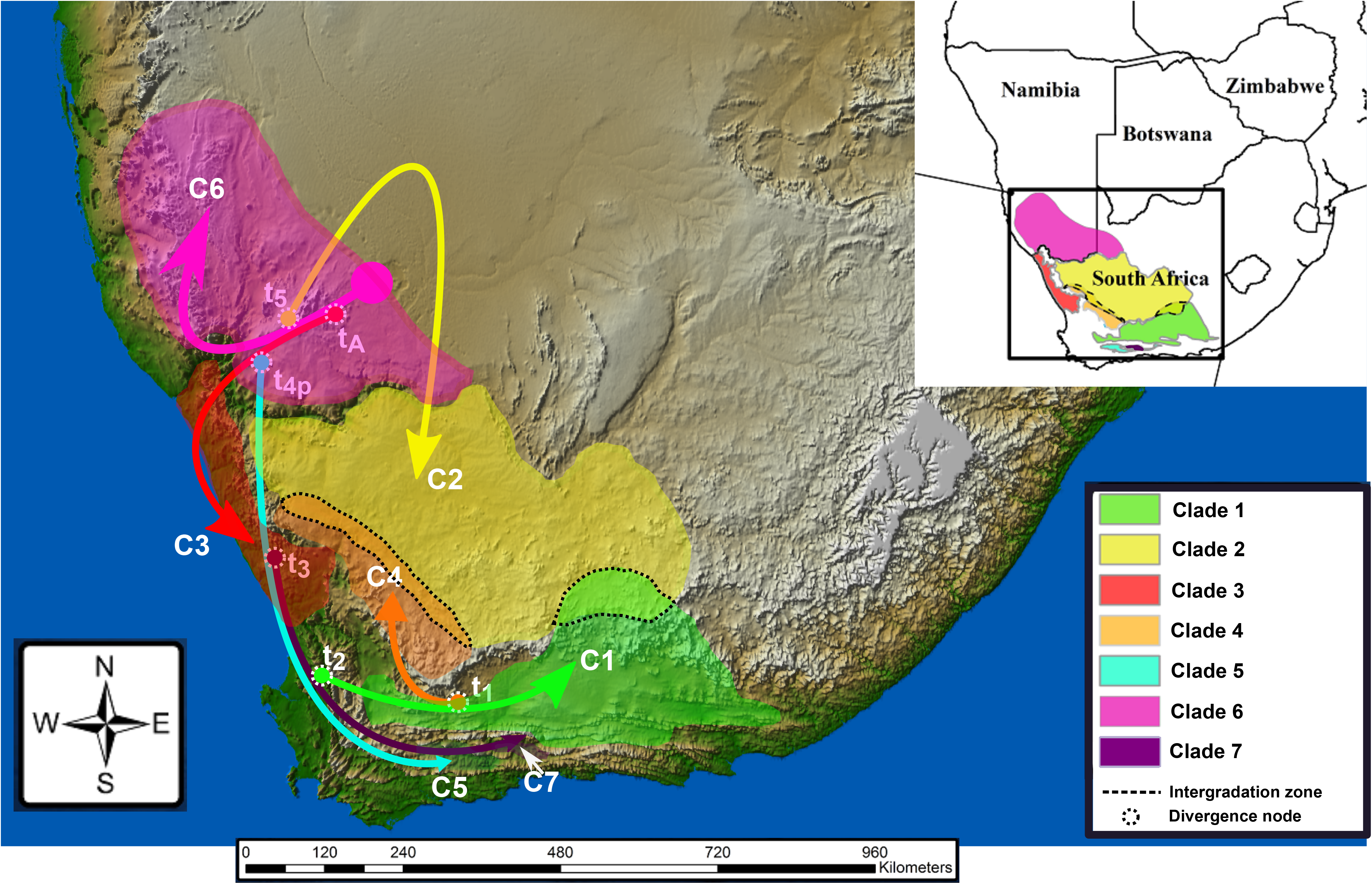
The most possible genealogy for the cladogenesis of the *Psammobates tentorius* complex suggested by the ABC simulation analysis based on total evidence datasets (mtDNA, nDNA and microsatellite DNA). Each critical divergence node that resulted from divergence events were indicated as circles. The arrows indicate the directional cladogenic process among the seven mtDNA clades.

### Morphological analyses

The Shapiro-Wilks test results and histograms indicated that log-transforming the datasets had improved their distribution but that they still did not fully conform to the requirements of a normal distribution.

### Sexual dimorphism

Two-way PERMANOVA results revealed significant morphometric differences among clades (F = 10.54, *p* < 0.0001) and size dimorphism between the sexes (F = 21.426, *p* < 0.0001). It also showed a significant interaction between sex and clades (F = 12.651, *p* < 0.005). This result also implied that the direction of sexual dimorphism was not fully consistent across clades.

For the phenotype-based sexual dimorphism, the coded character-based Two-way PERMANOVA results revealed significant character differences among clades (F = 7.81, *p* < 0.0001) and between sexes (F = 2.86, *p* < 0.0001). No significant interactions between sex and clades were, however, observed (F = 14.35, *p* = 0.14).

### Distinguishing morphologically among clades

The one-way PERMANOVA results revealed significant overall differences among clades for both females (F = 8.304, *p* < 0.0005) and males (F = 8.264, *p* < 0.0005). In females the Bonferroni corrected post-hoc test showed significant pairwise differences among all clades (with *p* < 0.01 in all cases), whilst, in males it failed to distinguish C1 from C5 and C3 from C4 (*p* > 0.05, in both cases).

The PCA analysis of the female dataset generated PC1 and PC2 as the two major principle components contributing 69.7% of the total variance. It generally revealed clear separation among groups (C1+C4+C5+C7), C3, C2 and C6, and the outgroup *Psammobates oculifer* (Kuhl, 1820) (Fig. 7). Moreover, it showed a slight separation between C1 and C4. However, the female based PCA was still not informative enough to separate all seven clades. In males, the PCA analysis also retrieved PC1 and PC2 as the two major principle components which contributed 63.7% of the total variance. It showed full separation of groups (C1+C5), C3, C4, C2 and C6. Since we were unable to obtain any male specimens for C7, its placement remains unknown. The outgroup *P. oculifer* was again fully separated from the *P. tentorius* species complex (Fig. 7).

**Figure 7.**
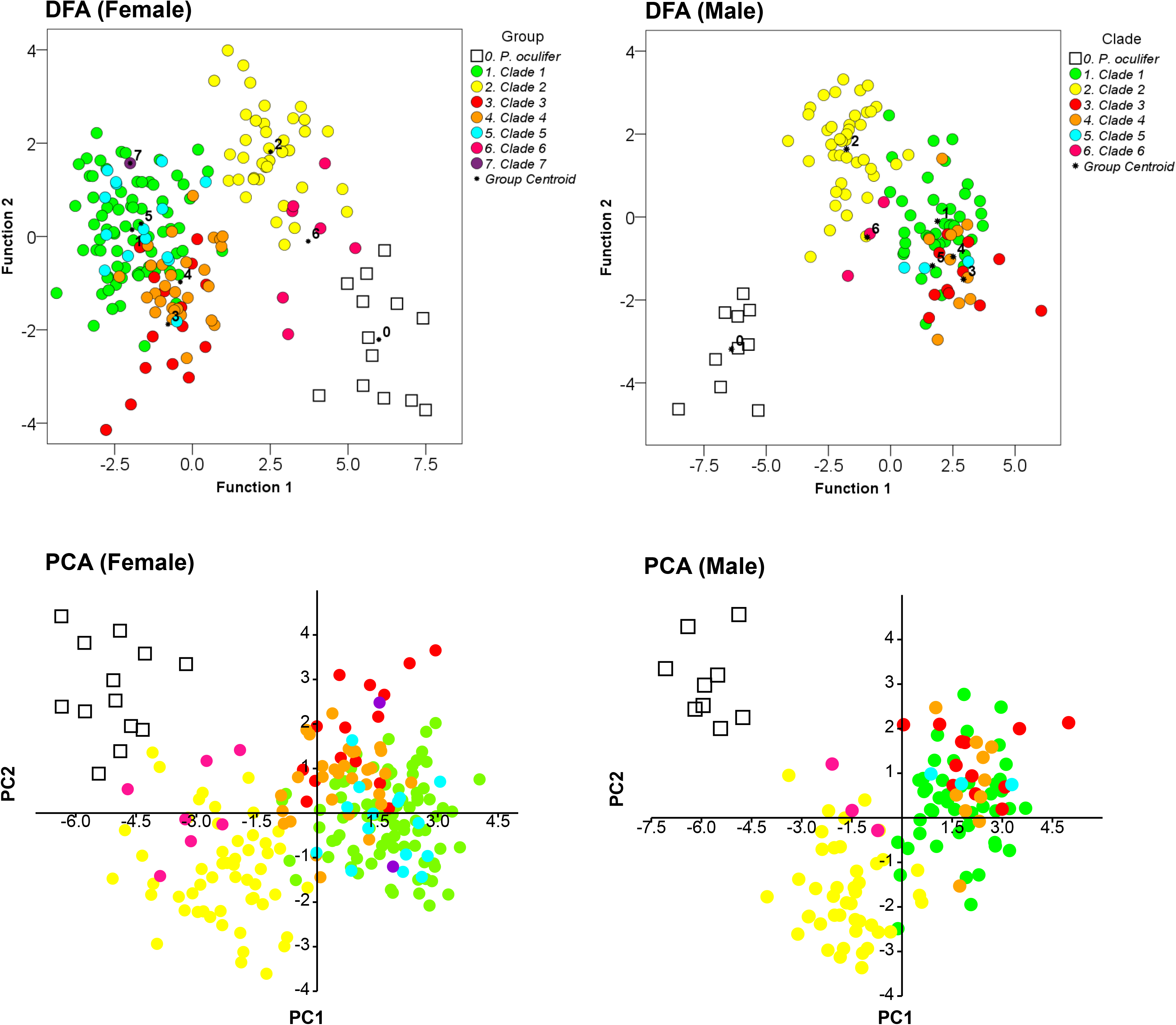
PCA and DFA scatterplots for the seven clades of the *P. tentorius* species complex across sexes (males and females), using the first two principal components and discriminant functions, respectively. The congeneric *P. oculifer* clearly stands apart from the clades of this study.

In the DFA analyses of females, the first two discriminant functions with the highest Eigen values contributed 81.83% of the total variance and significantly differentiated the seven clades from each other (Wilks’ lambda = 0.014, *p* < 0.0001). Overall, 87.18% of the cases were correctly assigned (C1: 89.61%, C2: 94.74%, C3: 86.67%, C4: 79.31%, C5: 53.33%, C6: 100%, C7: 100% and outgroup *P. oculifer*: 100%). The pairwise Mahalanobis distances revealed significant pairwise differences among the clades (*p* < 0.05 in all cases), except between C1 and C7 for which it was not significant (F = 1.12, *p* > 0.05). The DFA scatterplots (Fig. 7) generally exhibited four clusters: 1) C1+C4+C5+C7, 2) C3, 3) C2 and 4) C6. When comparing the pairwise Mahalanobis distances between the clades to their individual distances from *P. oculifer*, the interclade differences were smaller, showing their closer relatedness to each other than to *P. oculifer*. However, C1, C4, C5 and C7 did not show clear separation from each other.

As for the DFA in males, only six clades were included as no C7 samples were available. The first two discriminant functions with the highest Eigen values contributed 82.33% of the total variance and significantly differentiated the clades from each other (Wilks’ lambda = 0.007, *p* < 0.0001). In total 93.28% of the cases were correctly assigned (C1: 90.24%, C2: 100%, C3: 81.82%, C4: 80%, C5: 100%, C6: 100% and *P. oculifer*: 100%). The pairwise Mahalanobis distances revealed significant pairwise differences among the clades (*p* < 0.05 at all cases), except between C3 and C4, for which it was not significant (F = 0.882, *p* > 0.05). As in females, the DFA scatterplot of males also advocated four clusters. Within cluster 1, C1, C4 and C5 were not visually distinguishable.

The one-way PERMANOVA results of the female phenotypic data (see Supporting Information Table S13), also showed that the differences among the clades were significant (F = 21.172, *p* < 0.0001). The Bonferroni corrected post-hoc test results confirmed that all seven clades were significantly different from each other (*p* < 0.005 in all cases). In males, likewise, the one-way PERMANOVA results (see Supporting Information Table S13) revealed overall significant differences among the clades (F = 12.280, *p* < 0.0001). The Bonferroni post-hoc test indicated that the difference between C1 and C5 was not significant (*p* > 0.05), whilst, the differences among the rest of the clades were significant (*p* < 0.01 in all cases).

## DISCUSSION

### Genetic structure

The microsatellite DNA based STRUCTURE and DAPC analyses generally advocated 4 candidate species (Fig. 3). Clear subdivision within C6 from the DAPC, SAMOVA analyses and the mtDNA based phylogeny, suggested that further radiation between C6 (western) and C6 (eastern) was likely, and could lead to further cladogenesis. The western section of the Great Escarpment forms a geographic barrier that separates western and eastern population of C6, which may facilitate further divergence. Our Migrate analysis revealed that the gene flow rate among the four relatively close clades in the southern group “C1+C4+C7+C5”, was comparatively low (Supporting Information Table S12). A possible reason for this could be that geographical barriers reduced the gene flow between these clades. In the southern group, each clade occurs in an isolated geographic region, with the Swartberg Mountain, Rooiberg Mountain and the Langeberg Mountain separating C1, C7 and C5, respectively. The western side of the Great Escarpment separates C4 from C1, C7 and C5. These geographic barriers may have functioned as major driver in shaping the rapid divergence of the southern group.

The phylogenetic congruence between the combined mtDNA+nDNA and the mtDNA dataset may be an artefact resulting from the fast evolving mtDNA markers (faster substitution rate) dominating that of the nDNA markers. A further factor in this regard may be the larger number of mtDNA than nDNA markers used in the analyses. Notwithstanding this, Fisher-Reid and Wiens (2011) found that the combined mtDNA+nDNA data analyses were not necessarily dominated by the mtDNA dataset because of its more variable sites. In this study, the target organism is slow evolving (Avise et al., 1992; Tollis et al., 2017). In recent rapid radiation events, particularly in cases where there is a short time span in their rapid radiation history (i.e. in dealing with species complexes and slow evolving organisms like tortoises), tree topology conflict is particularly common. The most common consequence of Incomplete lineage sorting is the incongruence between gene and species trees (Maddison, 1997; Shaw, 2002; Rubinoff & Holland, 2005; Baum & Smith, 2013; Wang et al., 2018). Rubinoff & Holland (2005) suggested that mtDNA markers were more suitable and reliable for inferring phylogeny than slow evolving nDNA loci in studies focusing on slow evolving groups. Furthermore, nDNA markers were often found to have poor substitution signals due to their slow evolutionary and substitution rate, which results in poorly resolved phylogenetic tree topologies with poorly supported nodes in many studies (Zhao et al., 2020; Busschau et al., 2019), particularly in turtles (Caccone et al., 2004; Kindler et al., 2012; Petzold et al., 2014).

The shortcomings of pure mtDNA based phylogenetic inferences have also been addressed (Shaw, 2002; Hurst et al., 2005; Rubinoff & Holland, 2005; Wiens et al., 2010; Leliaert et al., 2014). These pitfalls may result in overestimations of species richness, misleading estimations of speciation rates and robustness of branch support, and incorrectly assigning the phylogenetic relationships between taxa. We should, however, remember that the nuclear genome and the mitochondrial genome are two separate regulatory entities, each retaining different evolutionary information, but both being crucial functional units of evolution (Rubinoff & Holland, 2005).

### Alternative cladogenic hypotheses

The ABC analyses did not retrieve results congruent with that of our phylogenetic analyses. Instead, it suggested C6 (basal lineage) as ancestral lineage and sister group to C2 (Fig. 6). In turn, C6+C2 was the sister group to C3. These results were not completely unexpected, and were in fact not unreasonable. The mtDNA based phylogenetic results gave the following groupings, (C2+C3)+C6 and (C1+C4)+C7)+C5, although there was uncertainty about the phylogenetic position of C6. The microsatellite based neighbour-joining tree suggested a sister relationship between C2 and C6. The rest of the topology was similar to that derived from the mtDNA dataset. The combined phylogeny was generally identical to that of the mtDNA phylogeny. The nDNA phylogeny also suggested that C2 and C6 were sister groups, and that C3 was more closely related to C1+C4+C5+C7. The best simulated scenario from the ABC analyses was indeed compatible with the results of all the datasets. The ABC simulated best scenario results of this study also suggested that there should be caution when solely using mtDNA data to infer a phylogeny. Morphologically, C3 was also more similar to the *P. t. tentorius* group (C1, C4, C5, C7), rather than the *P. t. verroxii* group (C2, C6) in terms of carapace shape and colour pattern. The information retrieved from the morphological characters also matched the findings of the ABC and nDNA analyses. Overall, the best scenario retrieved from the ABC simulation analyses was possibly the most likely one for explaining the real genealogy of *P. tentorius*, and it highlighted the importance of using multiple types of data to infer phylogenetic relationships among taxa.

### Does the Uniondale population represent a distinct clade?

The single 12S based phylogeny suggested that the Uniondale population was a close relative of C1, rather than C5 or C7 from the adjacent localities, which was not completely unexpected. Firstly, the geographic barrier between the southern distribution range of C1 and the Uniondale population is the eastern section of the Swartberg Mountain, which is much lower than its western section. The eastern section has ravines that may allow limited gene flow between the northern distribution range of C1 and the Uniondale population. Nonetheless, it is not clear whether the Uniondale population should only be considered as a subclade within C1, or as a distinct lineage and OTU, given the limited sequence data obtained from a single marker for museum specimens. When comparing the pairwise distances for the 12S gene, the genetic distance between the Uniondale population and C1 was smaller than the distance between C1 and C4. As our analyses generally suggested that C1 and C4 should not be regarded as different OTU’s or species, the Uniondale population should only be considered as a subclade of C1, rather than a distinct lineage or OTU.

The microsatellite DNA based pairwise difference matrix (Fig. 5), Slatkins linearized *F*_*ST*_ matrix (Fig. 5) and *F*_*ST*_ matrix (Supporting Information Figure S16) all advocated that C1 was closer to the Uniondale population, than to C5 or C7. In the relative divergence time tau matrix (Supporting Information Figure S16), it is clear that the estimated relative divergence time between C1 and the Uniondale population was the smallest of all, which again suggests that the Uniondale population is closest to C1. Further studies with fresh samples and more genes are therefore required to clarify its phylogenetic position.

### Hybridization and inbreeding

The microsatellite DNA based DAPC admixed individual analyses revealed possible hybridization between C1 and C2 at their intergradation zone (Victoria West and in the Richmond area). Similar findings of possible hybridization between C1 and C2 at these two localities were also made by Zhao et al. (2020). To further verify them as intergradation zones, additional extensive sampling is required. Although Zhao et al. (2020) also detected potential hybridization between C2 and C4, the present microsatellite DNA based analyses failed to detect any sign of crossing-over between C2 and C4. This may be due to the randomness of crossing-over that affects only part of the nuclear genome and may therefore have missed the microsatellite loci in the hybridized individuals.

Our Migrate analysis revealed potentially substantial gene flow in the two intergradation zones, between C1 and C2, and between C2 and C4 (see Supporting Information Table S12). In terms of direction, the gene flow rate was much higher from C1 to C2 than from C2 to C1. On the other hand, the gene flow rate from C2 to C4 was substantially higher than it was from C4 to C2. These findings implied possible hybridization between C1 and C2, and between C2 and C4, but this requires further verification. The Migrate results also suggested that the migrations in these intergradation zones were recent events. The divergence between C1 and C2 seems to have been influenced by the Great Escarpment (GE), since C1 mainly occur below the GE, whilst C2 is only present above the GE (Hofmeyr et al., 2014). The intergradation zone between C1 and C2 is in the middle section of the GE, where a clearly open corridor region seems to have recently allowed migration between C2 from the northern GE and C1 from the southern GE. Regarding the intergradation zone between C2 and C4, it seems that the biome was the major driver causing the divergence between C2 and C4, since C2 mainly occurs in the Nama Karoo, whilst C4 only occurs in Fynbos and Succulent Karoo (Hofmeyr et al., 2014). These assumptions, nonetheless require further verification.

Among the 11 populations, the C1 eastern population revealed extensive inbreeding. It occurs in the Fish River Valley and is isolated from other C1 populations. The Fish River Valley is isolated from the Nama Karoo habitat by Albany Thicket along the Fish River canyon (Mucina & Rutherford, 2011), resulting in the eastern population being isolated from central and western populations. Inbreeding may also have resulted from a bottleneck due to a substantial reduction in size of the Fish River Valley population because of habitat fragmentation. This was confirmed by the Arlequin analyses (Table 1) and suggests that the Fish River valley population may deserve special conservation attention. The strong inbreeding signal also detected in the C2 southern population, which occurs in a widely open region, is however unclear. This population occurs in the Nama Karoo, and is therefore possibly influenced by aridity and drought which may fragment the population, resulting in substantially reduced gene flow between subpopulations.

### Morphology

Efforts to develop an accurate and stable dichotomous key for the seven clades of the *P. tentorius* species complex was jeopardized by too much variations in potentially useful morphological diagnostic characters. The multivariate analyses generally advocated four highly parsimonious clusters: cluster 1) C1+C4+C5+C7, cluster 2) C3, cluster 3) C2 and cluster 4) C6, though some of the analyses advocated further subdivisions within cluster 1. The morphological clustering scheme revealed a good match with the nDNA sequence and microsatellite data clustering patterns. This supports the four-species scheme as the best taxonomic solution for the *P. tentorius* species complex (Fig. 4). When comparing the pairwise Mahalanobis distances between the seven clades, and between the seven clades and *P. oculifer*, the distances between the four morphological clusters were in some cases smaller than their distance from *P. oculifer*, but in other cases larger, specifically between clusters 1, 3 and 4. These morphological distances provided clear evidence that the clades were different at the interspecific rather than intraspecific level. However, given the relatively small sample sizes in the case of C5 and C7, but particularly in C7, verification of these findings with adequate samples is needed in their case.

The major morphological confusion in terms of both the carapace and plastron color and patterns came from C1, C2 and C6, on the one hand, and between C3 and C4 on the other hand (see Supporting Information Figures S1-S2). Clade 5 and C7 had far less within-group phenotypic morphological variations than the rest of the clades, but this may have been the result of their small sample sizes. Within C1, some individuals encountered in the Tankwa Karoo (south-eastern population) exhibited a uniformly brown carapace, which was also the case with some individuals in C2 (northern populations) and some in C6 (see Supporting Information Figures S1-S2). The placement of these uniformly brown carapace morphs was confirmed by both the DNA sequence data and microsatellite data. This occurrence could be the result of microhabitat adaptation and therefore convergent evolution, since these morphs were only found in habitats with sparse vegetation and barren sandy substrates (Z. Zhao, pers. obs.). Furthermore, some individuals from the central and south-western populations of C1 resembled similar phenotypes within C2 ((e.g. highly dull carapace patterns, and a faintly developed carapace dome and knobs – considered as typical characters of *P. t. verroxii* (Branch, 2008)). The molecular results confirmed that those individuals belonged to *P. t. tentorius* (C1), and not to *P. t. verroxii* (C2), which certainly contributes towards untangling its taxonomy. The microsatellite results revealed none of these strange “*P. t. verroxii* - like” individuals and thus confirmed that they were not a consequence of hybridization between putative taxa C1 and C2.

In the phenotypically equally confusing C2 (see Supporting Information Figures S1-S2), the northern and southern populations differed substantially. The molecular results (all markers), however, did not reveal any significant differences between the two populations. These results suggest that the phenotypic variations within C2 may have been shaped by short-term environmental adaptations, rather than long-term adaptive radiation. It is also noteworthy that many individuals encountered in the southern population were morphologically substantially similar to the typical phenotypes of C1 (e.g. highly domed carapace and distinct carapace stripes – considered as typical characters of *P. t. tentorius* by Branch, 2008). These confusing patterns would have resulted in substantial misidentifications between *P. t. verroxii* and *P. t. tentorius* in the past (see records of Hofmeyr et al., 2014), since many of the questionable individuals were found in the range of the southern population of C2. The microsatellite data failed to find any evidence to support these “*P. t. tentorius* – like cases” as resulting from the hybridization between individuals belonging to putative taxa C1 and C2, though both this study and Zhao et al. (2020) advocated possible crossing-over between C1 and C2.

As briefly mentioned by Zhao et al. (2020), the last part of the major confusion in the *P. tentorius* species complex is cause by the great phenotypic similarity between C3 and C4 (both previously regarded as “*P. t. trimeni*” by Boycott & Bourquin, 1988 and Branch, 2008), despite all the molecular evidence showing them to be distant relatives. The similarity could therefore be ascribed to homoplasy due to convergent evolution, as both occur in Succulent Karoo biomes (Branch, 1998 and 2008; Hofmeyr et al., 2014), thus most likely facing similar selection pressures.

### Taxonomic remarks

Species and OTUs are the fundamental entities of biological science (Blaxter et al., 2005; de Queiroz, 2007; Schmidt et al., 2014), and both are regarded as products and indicators of complex biotic and abiotic ecological interactions. Correctly determining OTUs is crucial in biological conservation and wildlife management (Avise, 1992; Moritz, 1994 & 2002; Thomson et al., 2018). The modern species concept is no longer restricted to morphology, but is rather linked to ecology with its temporal and spatial dimensions. Defining species or OTUs should therefore reflect evolutionary histories and phylogeographic processes (Avise, 1992; Moritz, 1994 & 2002), rather than empirical phenetics and morphology. Moreover, in the conservation of species, priority should be given to protecting important historical lineages and maximizing the preservation of speciation potential, rather than only focusing on endangered taxa (Avise, 1992; Moritz, 1994 & 2002). However, to correctly assign OTUs remains challenging (de Queiroz, 2007), the major challenges being a) difficulty in defining species boundaries (de Queiroz, 1998, 1999, 2005b, 2007) due to the absence of a unified species concept (Mallet, 2013), and b) unclarity about the criteria used to delineate species (de Queiroz, 1998, 1999, 2005a, b).

Besides the above issues, developing a proper OTU scheme for conservation management can be tough for a species complex when cryptic species are encountered (Heath et al., 2008). Ignoring the importance of correctly identifying cryptic species can lead to ineffective and costly conservation management (Briggs, 2005; Heath et al., 2008). In this study of the *P. tentorius* species complex, we discovered several cases of species crypsis with both the microsatellite and DNA sequence datasets, which would not have been possible without the use of these multiple molecular genetic markers. This was confirmed by the following findings: 1) most members of the southern population of C2 were previously regarded as *P. t. tentorius* based on morphological characters (Loveridge & Williams, 1957; Hofmeyr et al., 2014; Branch, 1998 & 2008), but were found to actually belong to *P. t. verroxii*; 2) despite being morphologically very different, the C2 southern and northern populations were found to not be genetically significantly different; 3) Loveridge & Williams (1957) and Boycott & Bourquin (1988) proposed that the southwestern Karoo (near Matjiesfontein and the Tankwa Karoo) as a possible “three-way contact zone” where the three subspecies coexist but the results confirmed that all individuals found in that region belonged to C1, despite phenotypically being a highly polymorphic population; 4) Loveridge & Williams (1957) and Branch (2008) suggested that the uniformly brown “*P. bergeri*” may deserve recognition as a valid taxon but the results showed that these uniformly brown morphs were present in C1, C2 and C6, and should therefore not be considered a valid taxon, but rather ecomorphs; 5) Boycott & Bourquin (1988) and Branch (2008) regarded the tortoises found above the Great Escarpment in the Kamiesberg, Hantam Karoo and Roggeveldberge region as *P. t. trimeni*, but the results confirmed the findings of Hofmeyr et al. (2014) and Zhao et al. (2020) that these tortoises belong to C4, the closest relative of C1, and not C3 (regarded as *P. t. trimeni*); true *P. t. trimeni* occurs only in the west coast succulent Karoo area; 6) although morphologically similar, C2 and C6 (both previously regarded as *P. t. verroxii*) are genetically significantly different; 7) morphologically C1 and C4 are distinguishable, but this study genetically recommends that they not be regarded as different taxa.

Based on all the genetic and morphological findings, and the multiple species delimitation results of Zhao et al. (2020), we recommend that at least (C1+C4+C5+C7), C2, C3 and C6 be recognized as full species (see Fig. 4). Fresh tissue samples are needed to determine the taxonomic status of the Uniondale population, and whether it represents a distinct taxon, or belongs to “C1+C4+C5+C7”. Detailed morphological descriptions still have to be carried out for these proposed new taxa and proper nomenclature developed based on the type specimens. Future studies of this complex should ideally be based on genome wide DNA data in order to unmask its full evolutionary history, and also to investigate the relationship between the genome and the extraordinary colour pattern variations by looking at gene expression and epigenetics.

## CONCLUSION

In terms of clustering scheme, the findings based on mtDNA, nDNA, microsatellite markers and morphology generally all supported the “four-species” hypothesis as the most parsimonious solution for solving the taxonomy of the species complex. The “four-species” were: 1) C1+C4+C7+C5, 2) C3, 3) C2 and 4) C6. The findings all indicated no significant difference between C1, C4, C7 and C5. The most likely scenario explaining the genealogy of the *P. tentorius* species complex was retrieved by the ABC simulation analysis. It suggested C6 as ancestral lineage and sister group to C2, and C6+C2 the sister group to C3. These findings were, however, not congruent with that of the phylogenetic analysis of Zhao et al. (2020), the later authors found C6 was the sister group to “C2+C3”. Signs of substantial inbreeding in the eastern population of C1 and southern population of C2 were found, caused either by their isolation from other populations or by a bottle-neck. This may have implications for their conservation. A hybridization signal was detected between C1 and C2, suggesting that crossing-over occurs between putative species C1 and C2.

## ACKNOWLEDGEMENTS

We thank the National Research Foundation (Grant - IFR150216114248) and the Universities of the Free State (UFS Research Grant: A1999/158110) and Western Cape (SNS Grants) for providing the funding to carry out the study. We also thank the following provincial departments of Nature Conservation for providing collecting permits: Northern Cape Province (Permits No: FAUNA 1061/2/2015, FAUNA 1266/2016, FAUNA 1458/2015, FAUNA 1267/2016, FAUNA 0729/2018, FAUNA 0730/2018), Western Cape Province (Permit No: AAA007-00179-0056) and Eastern Cape Province (Permits No: CRO 171/16CR, CRO 172/16CR), as well as the Ministry of Environment and Tourism, Namibia (Permit No: 1430/2009). The Animal Research Ethics Committees of the University of the Free State (AREC References No: 180111-005, UFS-AED2015/0013) and University of the Western Cape (ScRiRC2008/39) are thanked for providing ethical clearance for the research. The High-Performance Computing Centre of the University of the Free State is thanked for assisting with the phylogenetic analyses.

## SUPPORTING INFORMATION

### Figures

Figure S1. Morphs in the *Psammobates tentorius* species complex across the seven mtDNA clades, showing the high level of carapace phenotypic variation. The phylogenetic trees are modified versions of the ones published by Zhao et al. (2020). Left side: the phylogenetic relationships across the seven mtDNA clades; right side: the phylogenetic relationships among the seven clades based on nDNA data. The uniformly brown “*Psammobates bergeri*” morph was placed in red boxes; individuals which look like “*P. t. trimeni*” but belong to Clade 4 (a *P. t. tentorius* clade) were placed in orange boxes; individuals from Clade 2 (*P. t. verroxii*, south of the Orange River, Upper Karoo region) which look like “*P. t. tentorius*” were placed in green boxes; and individuals assigned to Clade 1, but which look like “*P. t. verroxii*, were placed in yellow boxes. The configuration of the phylogenetic tree showing the “three-subspecies” assumption is indicated on the right side.

Figure S2. Phenotypic variation in plastron patterns in the seven clades of the *Psammobates tentorius* species complex. The mtDNA clades (left side) and nDNA clades showing the currently recognised “three-subspecies” assumption (right side) are modified versions of the phylogenetic tree in Zhao et al. (2020).

Figure S3. Measurements of the plastron and carapace. P1: (1) CLS, (2) UW, (3) CW and (4) LW. P2: (1) CLS and (2) CLE. P3: HL. P4: (1) PLE, (2) AILI and (3) AILO.

Figure S4. Measurements of the plastron. (1) PLS, (2) AWD, (3) AWT, (4) FWT, (5) FALS, (6) FALC, (7) FWD, (8) GPLC, (9) GPLS, (10) GWT, (11) GWD, (12) HWT, (13) HWD, (14) OHW. (15) OPW and (16) OAW.

Figure S5. Results of BIC value versus the number of clusters to determine the optimal clustering scheme in the DAPC analyses. The a-score optimization determined the optimal number of retained principal components (PCs) and the proportion of successful predictions versus number of PCA axes retained. The Cross-Validation analysis determined the initial number of PCs retained.

Figure S6. Allele diversities among different localities (only localities with n ≥ 3 are shown) for the 14 different microsatellite DNA loci. The numbers on the x-axis represent different loci.

Figure S7. Membership prediction plot for ten suspected admixed individuals to determine potential hybridizations. “C1_E”: C1 (eastern), “C1_W”: C1 (western), “C2_N”: C2 (northern), “C2_S”: C2 (southern), “C6_E”: C6 (eastern) and “C6_W”: C6 (western).

Figure S8. Histograms of inbreeding scores among different populations. Groups with frequency bars above 0.4 were considered as showing high inbreeding levels.

Figure S9. The single marker (12S) based tree retrieved from the BI and ML analyses. The tree topology was derived from a ML analysis. Support values are given for BI (left) and ML (right); “*” indicates strong support (BP > 70 or PP > 0.95).

Figure S10. The combined (mtDNA+nDNA) based ML tree; “*” indicates weak support (BP < 70); all nodes without “*” were strongly supported (BP > 70).

Figure S11. A: The time calibrated species tree inferred by Bayesian analyses with the StarBeast package in BEAST 2; B: The time calibrated gene tree generated by the StarBeast analysis; the seven red dots represent the seven constrained calibration nodes. On both the gene tree and species tree, the blue bar represents the estimated 95% HPD time range; the estimated age at each node is shown in the middle of each 95% HPD bar.

Figure S12. Best model selection using model comparison methods implemented in the ABC analyses. The scenario with highest Direct score (occupying the highest proportion) was considered the most plausible scenario.

Figure S13. Scatterplot showing the PCA results evaluating the scenarios and priors in the ABC analyses. Each small dot represents a simulated dataset from the reference table and the large yellow dot represents the observed dataset. I expected the yellow dot to fall in the range of the small dots, which indicated that sampling was adequate.

Figure S14. Results of the model checking ABC analyses. The observed dataset (shown as the big yellow dot) fell within the range of both the simulated priors (open dots) and posteriors (medium size dots) datasets, showing that sampling was adequate, and the simulation analysis results reliable.

Figure S15. The parameters involved in the ABC analyses between priors and posteriors. The majority of the distributions of the priors and posteriors revealed a good match, implying that the simulation analysis results were reliable.

Figure S16. The relative divergence time (tau) matrix and the pairwise *F*_*ST*_ matrix generated among clades from the AMOVA analyses between mtDNA (shown on the left side) and microsatellite DNA (shown on the right side) datasets. “UN” refers to the Uniondale population.

### Tables

Table S1. List of all samples, their corresponding localities and NCBI GenBank accession numbers across different genes.

Table S2. Primers used in the study with corresponding oligo sequences, optimized annealing temperatures and sources.

Table S3. Allele size ranges, repeat motifs with their NCBI GenBank accession numbers, Multiply-mix reaction grouping schemes, the oligo-nucleotide sequences of primers with dyes, optimal annealing temperatures and the sources of primers of all microsatellite DNA markers tested in this study.

Table S4: Optimal partition scheme, substitution model, likelihood score (-InL), Gamma shape, proportion of estimated invariant and Homogeneity Test results.

Table S5. GenBank accession numbers for all outgroups used in the study. Also, the seven calibration nodes with their mean and range as used in the study with literature referenced.

Table S6. The list of all morphometric measurements and ratios used in this study, together with detailed information on specimens, clade belonging, location collected, country, sex and life stage. “F” = female, “M” = male. “PEM R” represents specimens deposited in Port Elizabeth Museum, “ZPT” represents specimens measured from this study, “ZR” represents specimens stored in Cape Town Iziko Museum.

Table S7. The list of all phenotypic characters used in this study, together with detailed information on specimens and their clade, locality collected, country, sexes and life stage. “F” = female, “M” = male. “A”= Adult, “J” = juvenile. “PEM R”, specimens from Port Elizabeth Museum, “ZPT”, specimens measured in this study, “ZR”, specimens from Cape Town Iziko Museum, “SMN”, specimens stored in Namibia National Museum. All other ID codes of specimens represent local municipal museums.

Table S8. The genotyping results across all 19 microsatellite DNA loci.

Table S9. The genotyping results of 14 microsatellite DNA loci used in this study, together with locality and subpopulation information.

Table S10. SAMOVA analysis results for the mtDNA and microsatellite DNA datasets across all scenarios (tests with different cluster schemes from K=2 to K=12). “*” indicates p < 0.05, “**”, p < 0.01 and “***” p < 0.001

Table S11. The genetic diversity analysis results of Arlequin showing locality information, sample size, geographic coordinates, polymorphic sites, number of haplotypes (h), haplotype diversity (Hd), average number of nucleotide differences (K), nucleotide diversity (Pi), gene diversity, genetic diversity (Theta, H), polymorphic loci and number of alleles (Allele No.) among different localities, using the mtDNA and microsatellite DNA datasets.

Table S12. The Migrate analysis results (using microsatellite data) estimated effective population sizes of the seven mtDNA clades (Θ1-Θ7), and the potential gene flow rate (M). M1->2 denotes the gene flow rate from Clade 1 to Clade 2. The directional gene flows occurring in the two intergradation zones, between Clade 1 and Clade 2, and between C2 and C4 are in bold.

Table S13. Summary of results retrieved with multivariate approaches, PERMANOVA, DFA with continuous and discrete datasets, with sexes analyzed independently to determine the most likely clustering schemes.

