## Supporting Information Document for "Testing alternative phylogenetic hypotheses for the tent tortoise species complex (Reptilia, Testudinidae) using multiple data types and methods"

*Morphological analyses*

*Specimen and data collection*

In order to obtain adequate datasets for the phenotypic characters (discrete data) and morphometric measurements (continuous data), specimens were collected in the field as well as obtained from the following museums, Bayworld Museum (Port Elizabeth), Iziko Museum (Cape Town), the Ditsong Museum (Pretoria) and the National Museum of Namibia (Windhoek). Specimens were carefully selected, and individuals with visibly abnormal character states were excluded from the analyses. All specimens were labelled and photographed dorsally, ventrally, anteriorly, posteriorly and laterally. In terms of wild caught specimens, a permanent marker was used to mark specimens prior to their release at the site of capture, in order to avoid repeat sampling. As for the continuous data, measurements were taken of the different linear aspects of the carapace and plastron using an electronic Vernier caliper and a tape measure. Each variable of interest was measured twice, and the average used in order to increase accuracy. In terms of the morphometric analysis, a closely related congeneric species, *Psammobates oculifer* (Kuhl, 1820) (Hofmeyr et al., 2017), was included in the multivariate analyses to assess how the relative morphometric distances among the seven clades recognized with the DNA sequence data compared to their relative morphometric distances to *P. oculifer*.

For continuous dataset, A total of 349 specimens were used for the morphometric analyses, comprising 221 adult females, 128 adult males (see Supporting Information Table S6). Twenty adult *P. oculifer* specimens were included in the analyses for purposes of comparison. For the phenotypic analyses 313 specimens were used, which included 224 females, 89 (see Supporting Information Table S7).

*Grouping variables and data partitions*

The seven clades retrieved by Zhao et al. (2020) were used as grouping variables for the univariate and multivariate analyses of both the discrete and continuous datasets. The morphological data were partitioned into phenotypic and morphometric datasets. Males were regarded as adult if their straight-line plastron length (PLS) exceeded 55.0 mm and straight-line carapace length (CLS) exceeded 65.0 mm. Females were regarded as adult if their PLS exceeded 75.0 mm and CLS exceeded 85.0 mm.

*The morphometric datasets*

A preliminary univariate factorial ANCOVA was carried out to determine whether variables were informative and stable (i.e., consistently showed significant results for all the clades) within groups, using IBM^®^SPSS^®^ version 20, before they were included in the dataset. The CLS (carapace straight-line length) was used as covariate.

The following carapace and plastron measurements were taken, see Supplementary Figure S3-S4, CLS, UW (upper width of carapace), CW (central width of carapace), LW (lower width of carapace), CLE (exact midline carapace surface length), HL (carapace height length), PLE (exact plastron length), AILI (inner axillary inguinal length) and AILO (outer axillary inguinal length); for detail about measurements, see Supporting Information Table S6.

In Supplementary Figure S3-S4 the following is shown, (1) PLS (straight-line plastron length); (2) AWD (lower anal width); (3) AWT (upper anal width); (4) FWT (upper femoral width); (5) FALS (lateral femoral to anal length); (6) FALC (midline femoral to anal length); (7) FWD (posterior femoral width); (8) GPLC (midline gular to pectoral length); (9) GPLS (lateral gular to pectoral length); (10) GWT (anterior gular width); (11) GWD (posterior gular width); (12) HWT (upper humeral width); (13) HWD (humeral width at bottom); (14) OHW (humeral width outward); (15) OPW (pectoral width outward) and (16) OAW (Anal side width outward); for detail, see Supplementary Table S6.

*The phenotypic datasets*

The same criteria used for selecting the continuous variables were also applied in the case of the discrete variables. Characters were encoded for the multivariate PERMANOVA (Tatsuoka, 1988). Two categories of coding were applied: 1) if a character only had two states, namely, present or absent, it was encoded as “1”, for “present” and “2” for “absent”. Character states encoded as 1 included: radiated sparse PCP, unmarked plastron, cone stripe, apical ring X, apical ring knob, orange head skin, orange buttock, white head block, black head apical speckle, gular triangular mark and triple mark. 2) in those cases where characters had multiple states, the following coding was used: “absent”, encoded as “0”, “present at low level”, encoded as “1”, “present at low to moderate level”, encoded as “2”, “present at moderate level”, encoded as “3”, “present at moderate to high level”, encoded as “4”, “present at high level”, encoded as “5”, present at very high level”, encoded as “6”.

Characters encoded according to the category 2 system included: striped fork, dull striped, solid central plastron pattern (PCP), radiated dense PCP, plastron side stripe, thick stripe richness, thin stripe richness, increasing stripe richness, equal stripe richness, stripe general thickness, discontinued stripe level, head smoothness, apical ring size, apical ring spot, drab stripe, orange plastron, orange striped, red striped, red marginal, light phase, dark phase, yellowish head block, buttock tubercle, carapace smoothness, knob scute status, domed carapace, forelimb distinct border, shell thickness, bent stripes and stripe density. The summarized character dataset is given in Supplementary Table S7.

**Supplementary Figures**

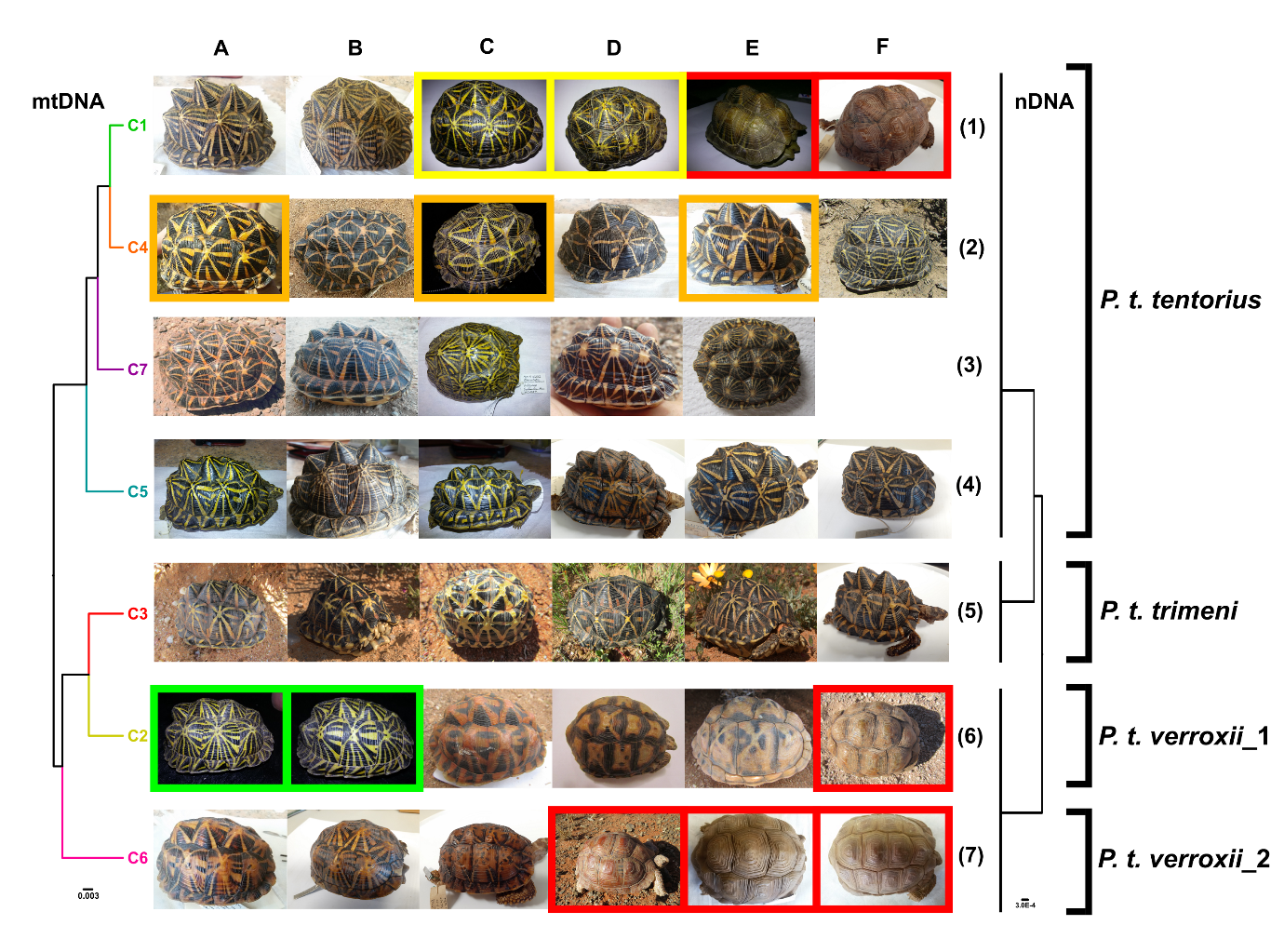

Figure S1. Morphs in the *Psammobates tentorius* species complex across the seven mtDNA clades, showing the high level of carapace phenotypic variation. The phylogenetic trees are modified versions of the ones published by Zhao et al. (2020). Left side: the phylogenetic relationships across the seven mtDNA clades; right side: the phylogenetic relationships among the seven clades based on nDNA data. The uniformly brown “*Psammobates bergeri*” morph was placed in red boxes; individuals which look like “*P. t. trimeni*” but belong to Clade 4 (a *P. t. tentorius* clade) were placed in orange boxes; individuals from Clade 2 (*P. t. verroxii*, south of the Orange River, Upper Karoo region) which look like “*P. t. tentorius*” were placed in green boxes; and individuals assigned to Clade 1, but which look like “*P. t. verroxii*, were placed in yellow boxes. The configuration of the phylogenetic tree showing the “three-subspecies” assumption is indicated on the right side.

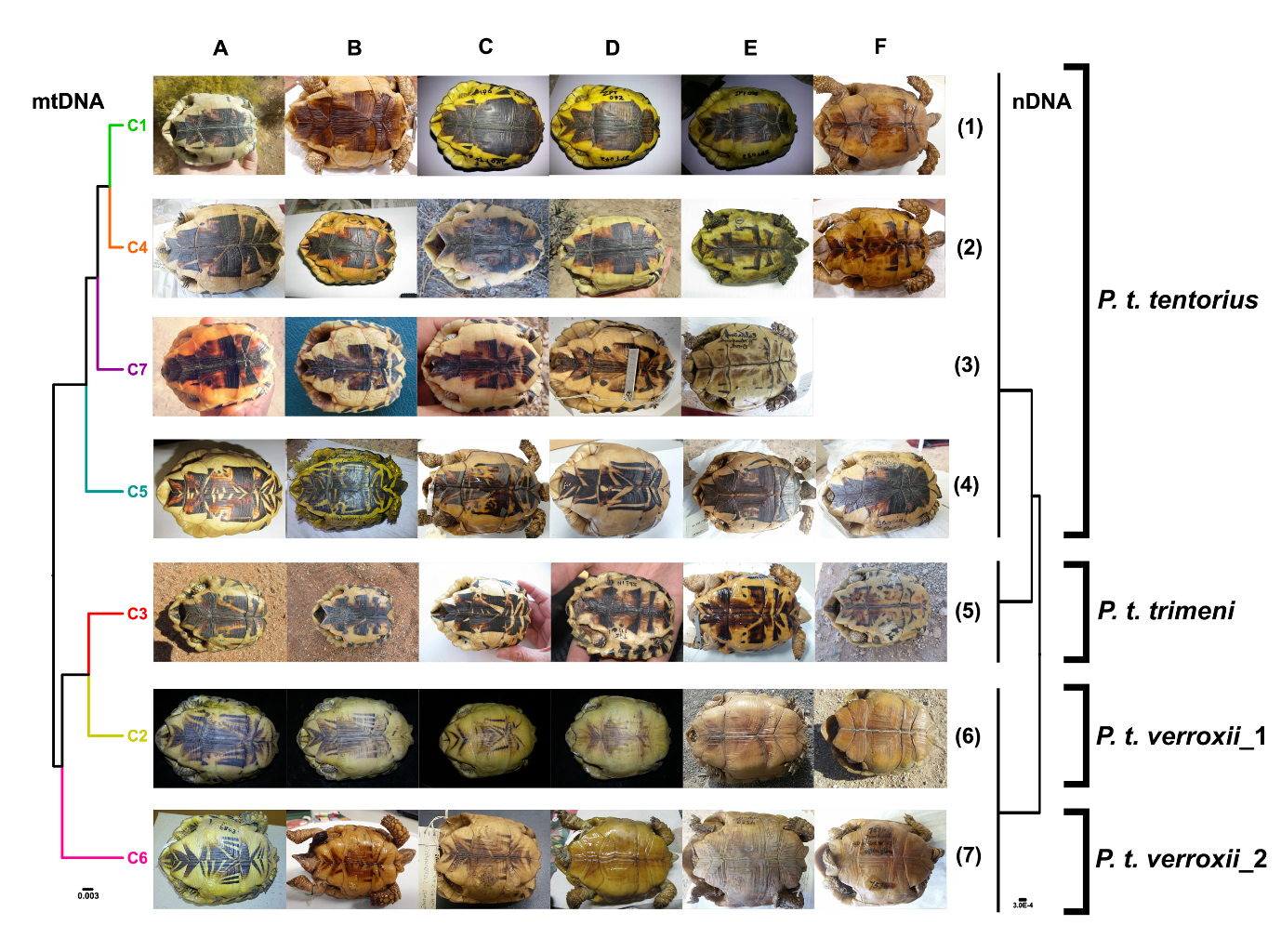

Figure S2. Phenotypic variation in plastron patterns in the seven clades of the *Psammobates tentorius* species complex. The mtDNA clades (left side) and nDNA clades showing the currently recognised “three-subspecies” assumption (right side) are modified versions of the phylogenetic tree in Zhao et al. (2020).

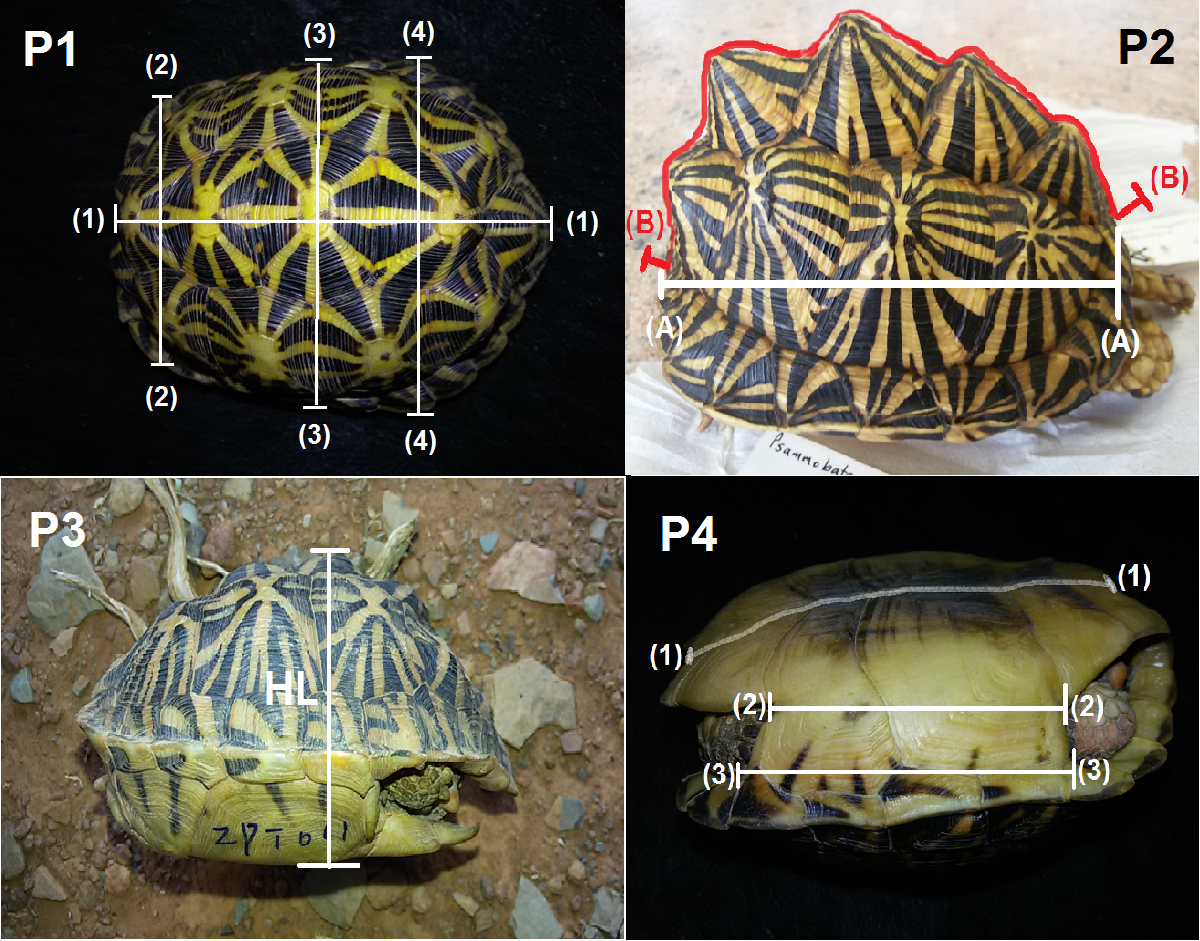

Figure S3. Measurements of the plastron and carapace. P1: (1) CLS, (2) UW, (3) CW and (4) LW. P2: (1) CLS and (2) CLE. P3: HL. P4: (1) PLE, (2) AILI and (3) AILO.

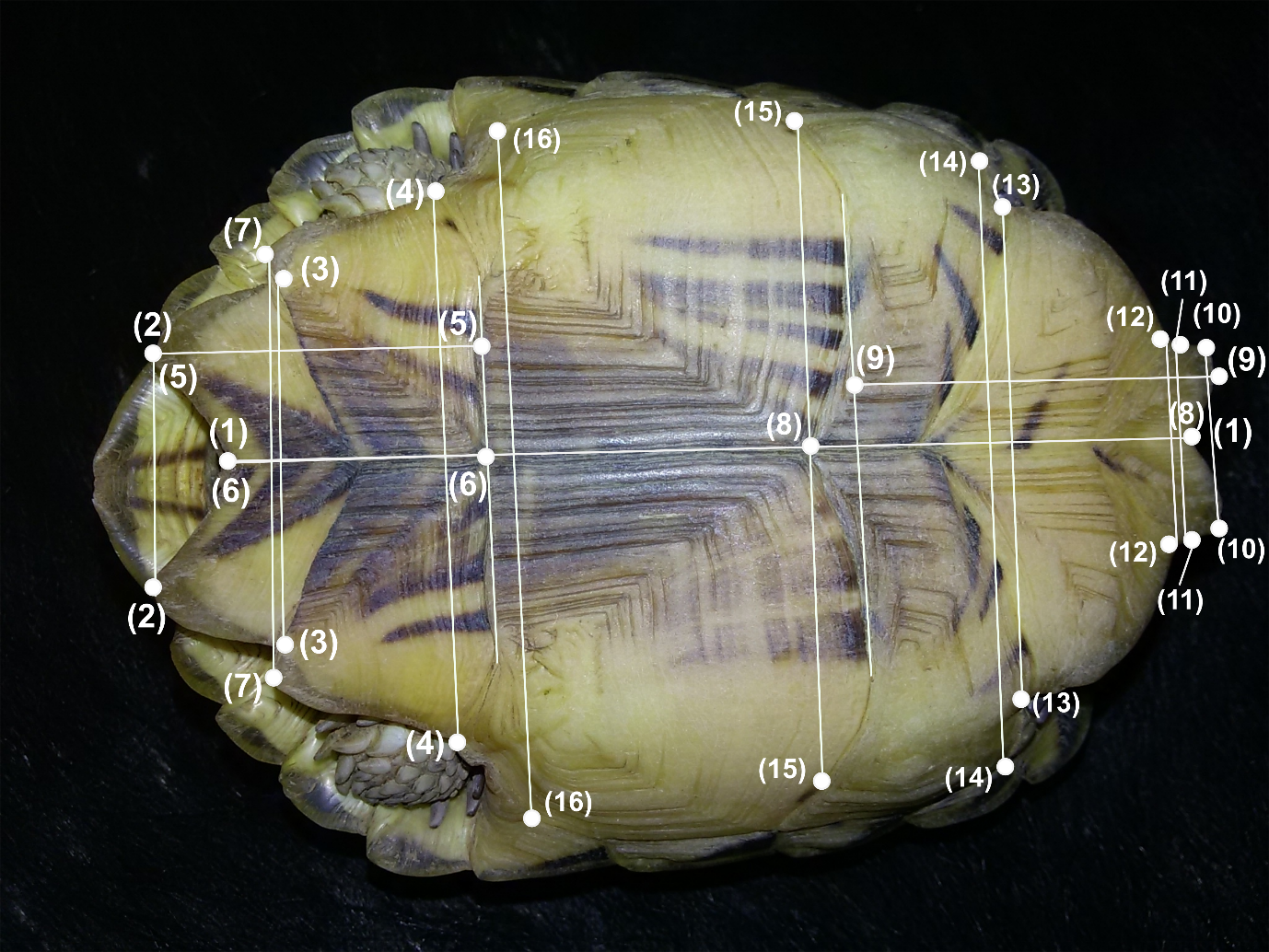

Figure S4. Measurements of the plastron. (1) PLS, (2) AWD, (3) AWT, (4) FWT, (5) FALS, (6) FALC, (7) FWD, (8) GPLC, (9) GPLS, (10) GWT, (11) GWD, (12) HWT, (13) HWD, (14) OHW. (15) OPW and (16) OAW.

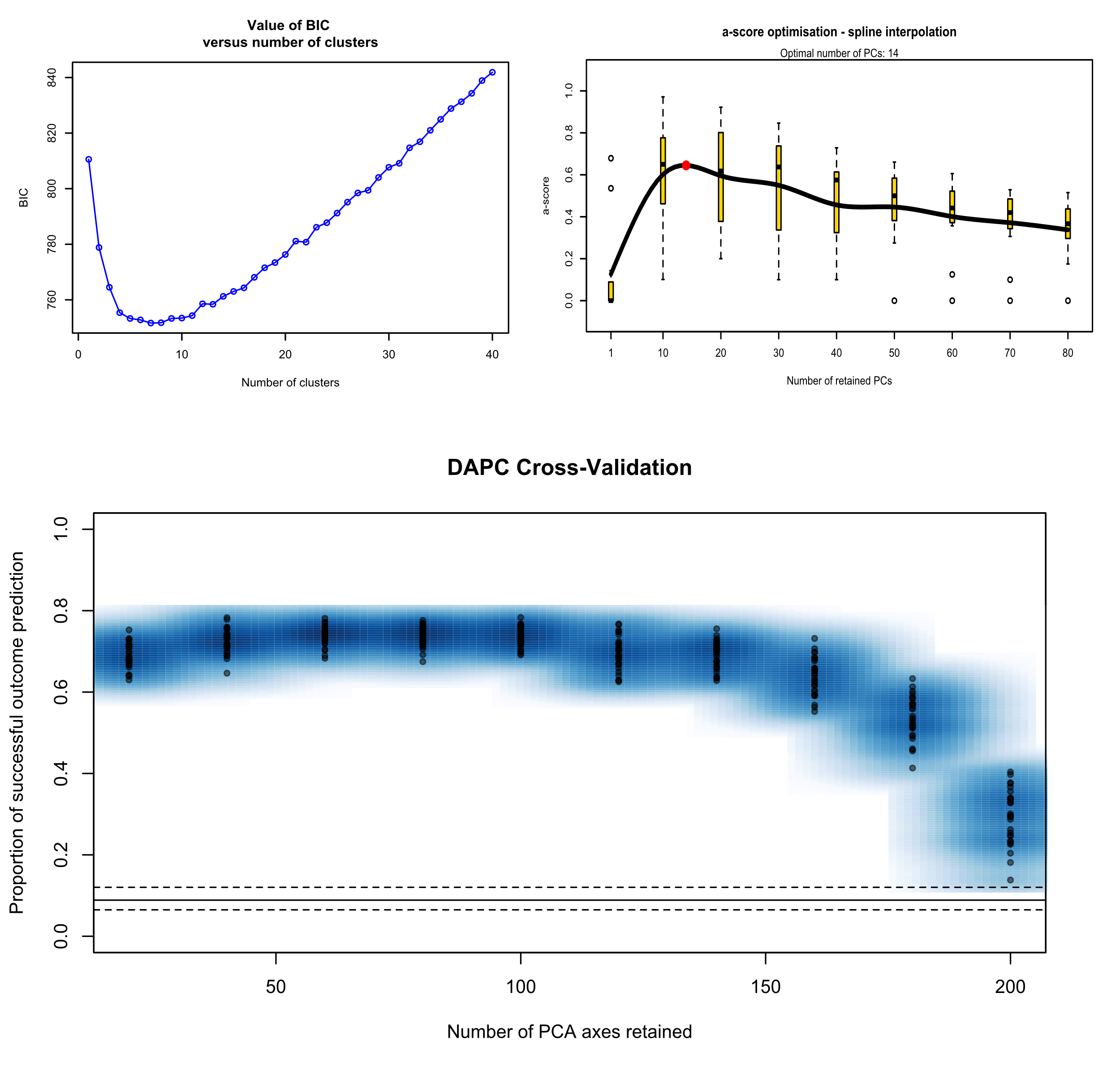

Figure S5. Results of BIC value versus the number of clusters to determine the optimal clustering scheme in the DAPC analyses. The a-score optimization determined the optimal number of retained principal components (PCs) and the proportion of successful predictions versus number of PCA axes retained. The Cross-Validation analysis determined the initial number of PCs retained.

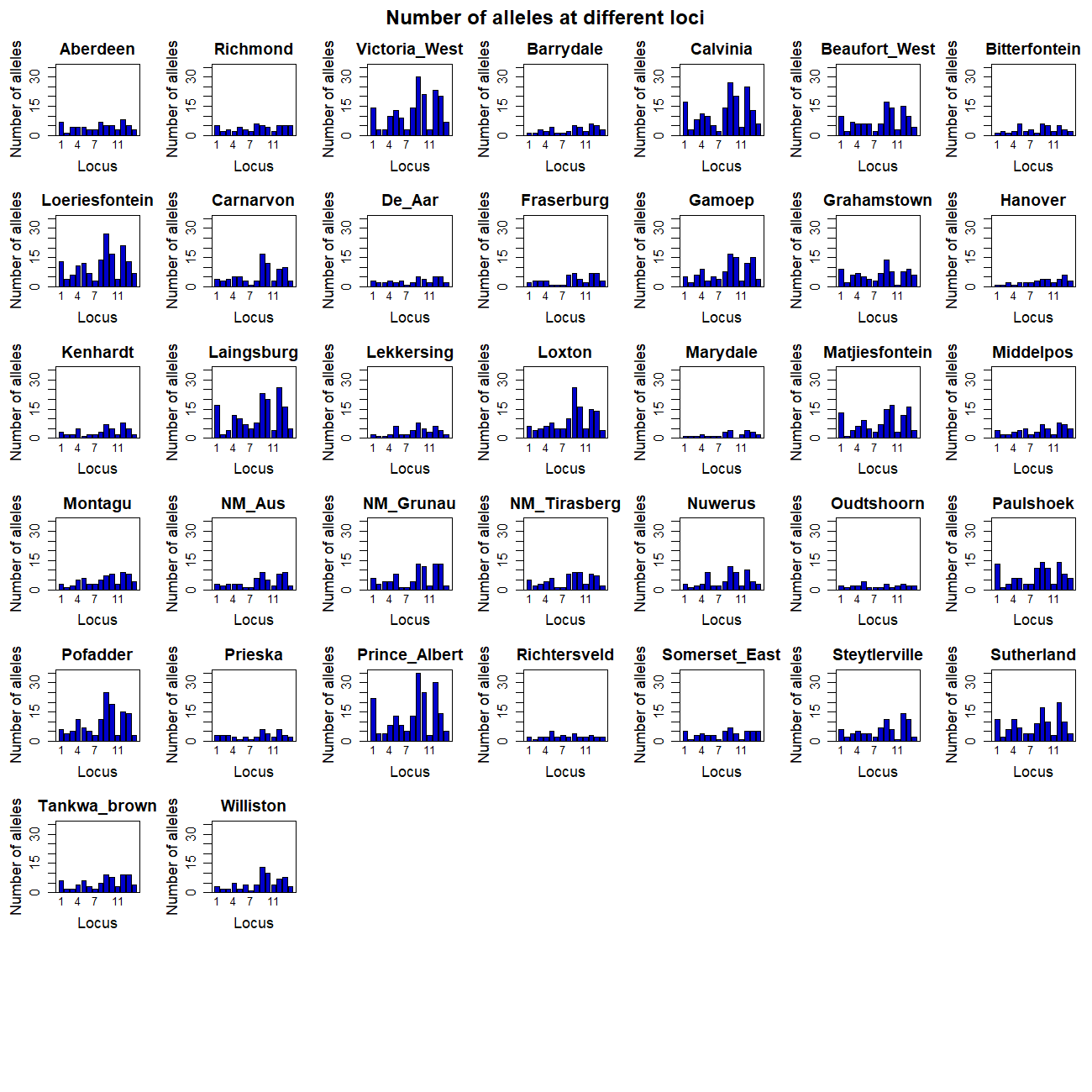

Figure S6. Allele diversities among different localities (only localities with n ≥ 3 are shown) for the 14 different microsatellite DNA loci. The numbers on the x-axis represent different loci.

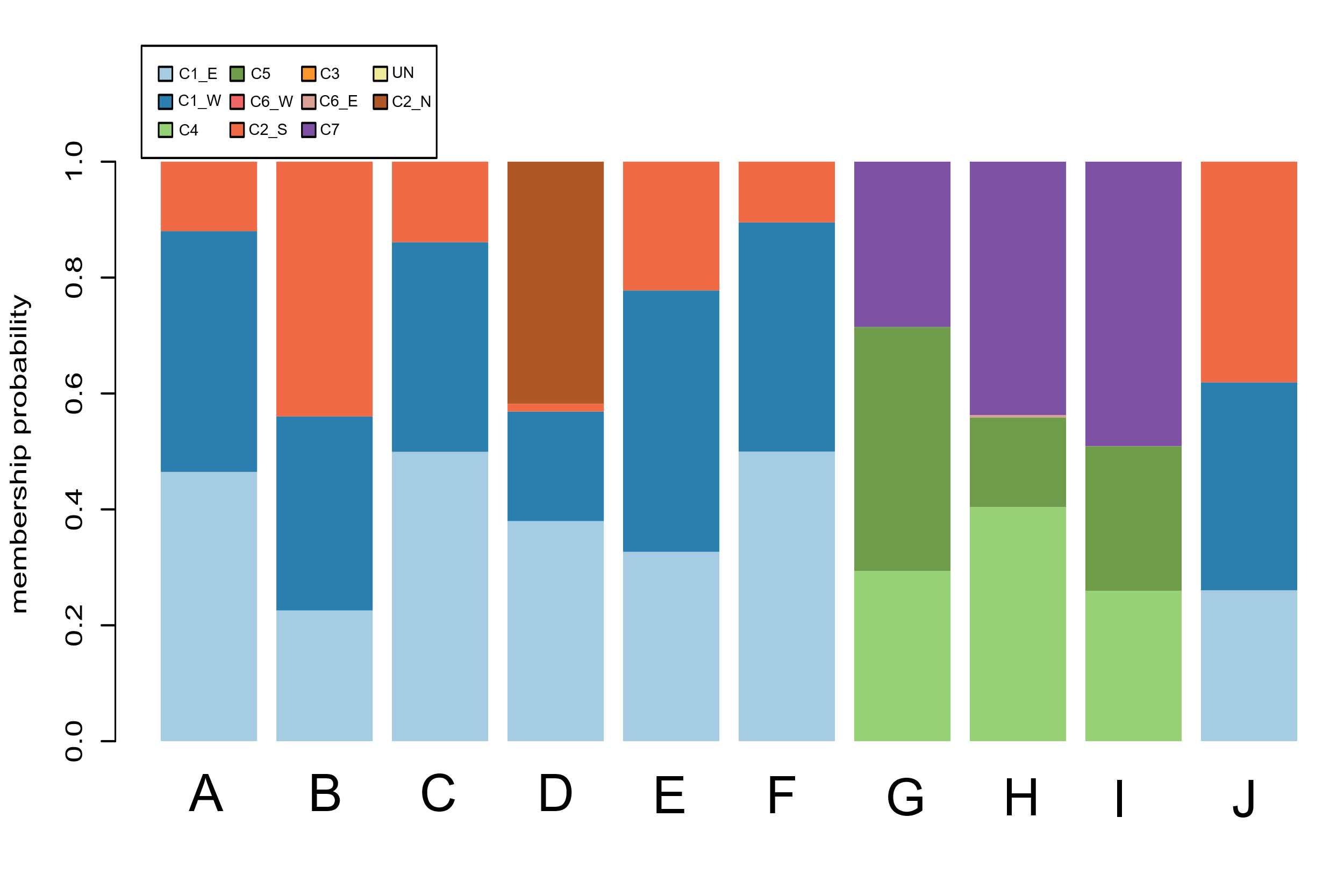

Figure S7. Membership prediction plot for ten suspected admixed individuals to determine potential hybridizations. “C1_E”: C1 (eastern), “C1_W”: C1 (western), “C2_N”: C2 (northern), “C2_S”: C2 (southern), “C6_E”: C6 (eastern) and “C6_W”: C6 (western).

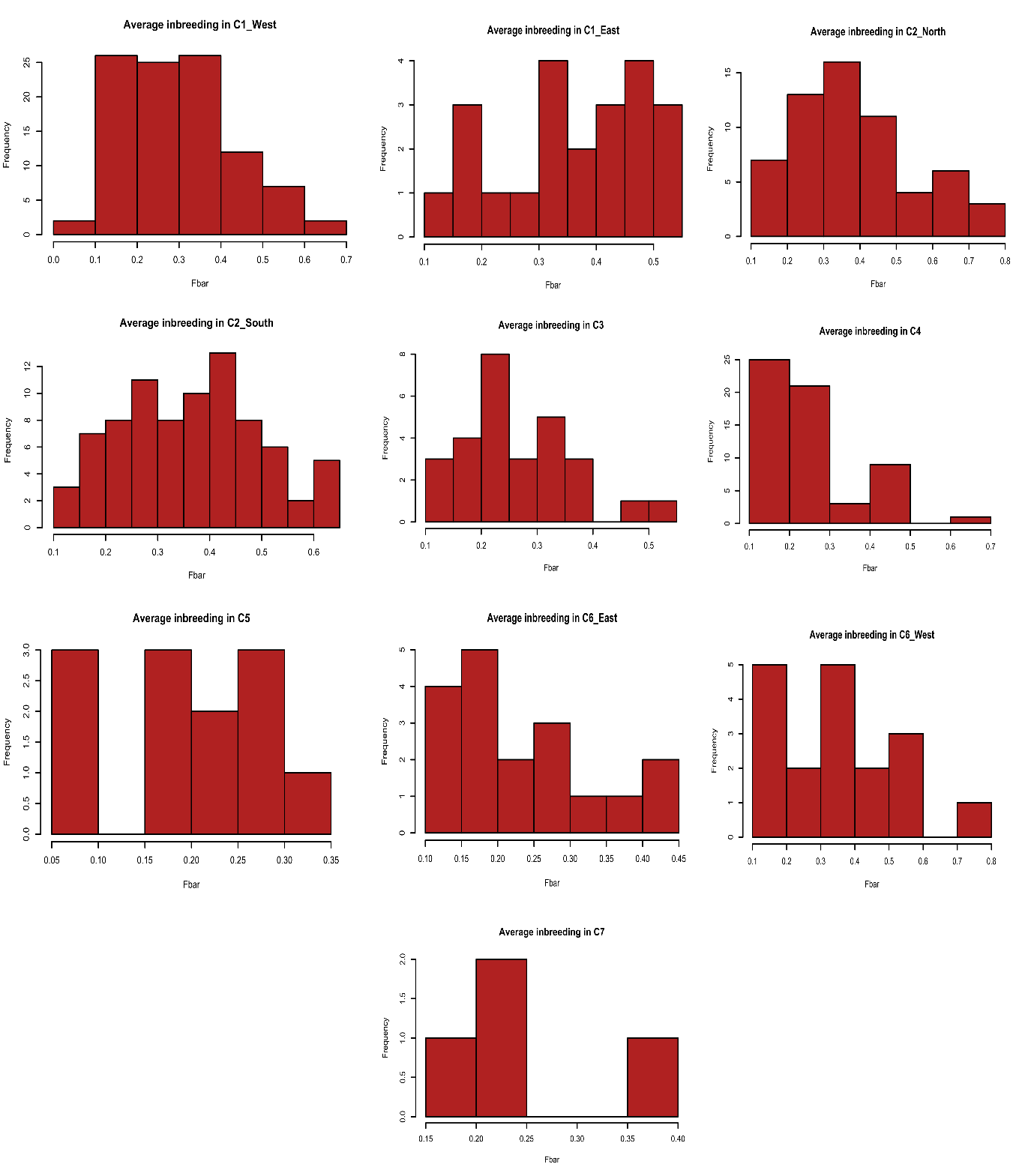

Figure S8. Histograms of inbreeding scores among different populations. Groups with frequency bars above 0.4 were considered as showing high inbreeding levels.

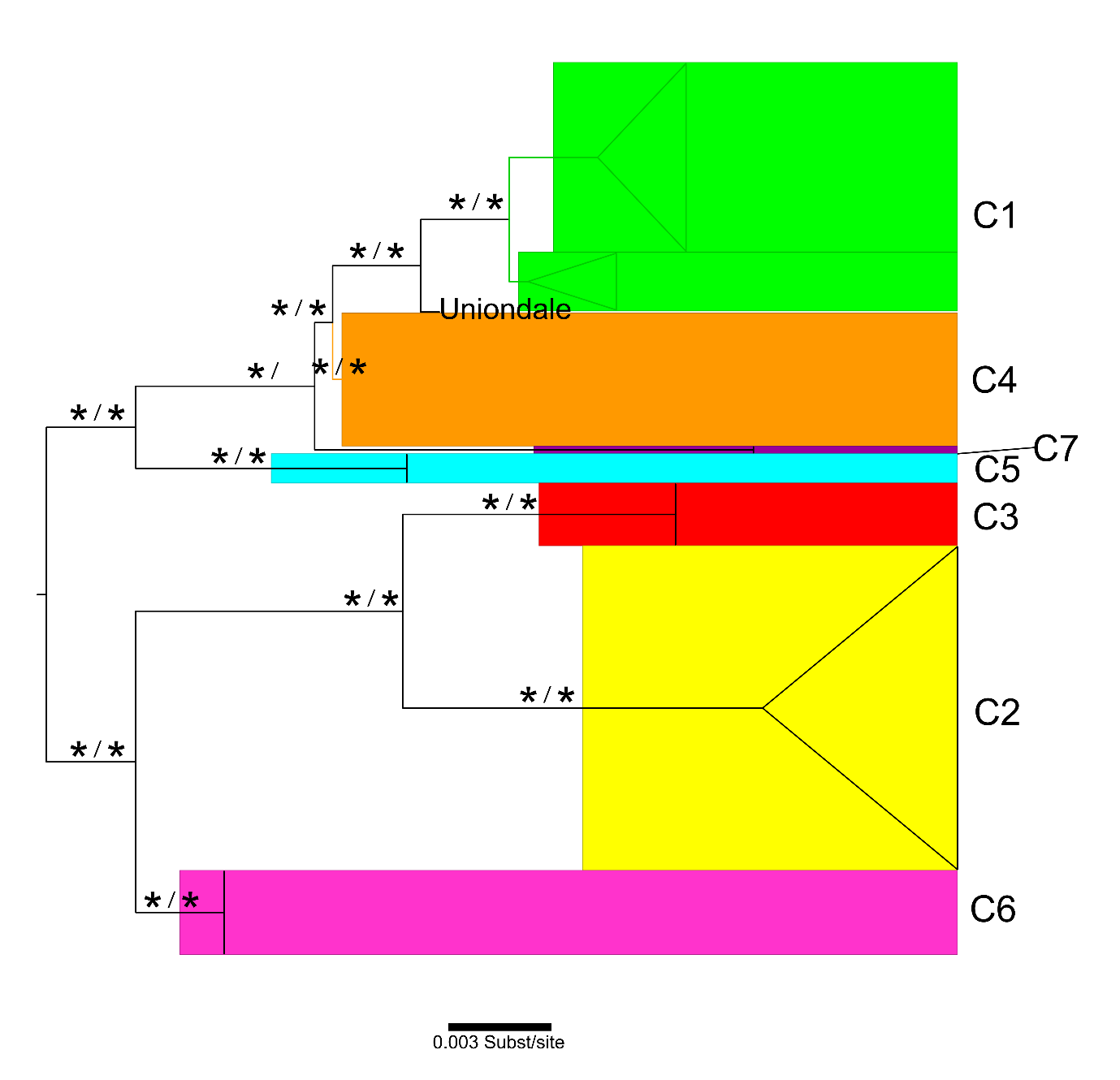

Figure S9. The single marker (12S) based tree retrieved from the BI and ML analyses. The tree topology was derived from a ML analysis. Support values are given for BI (left) and ML (right); “ * ” indicates strong support (BP > 70 or PP > 0.95).

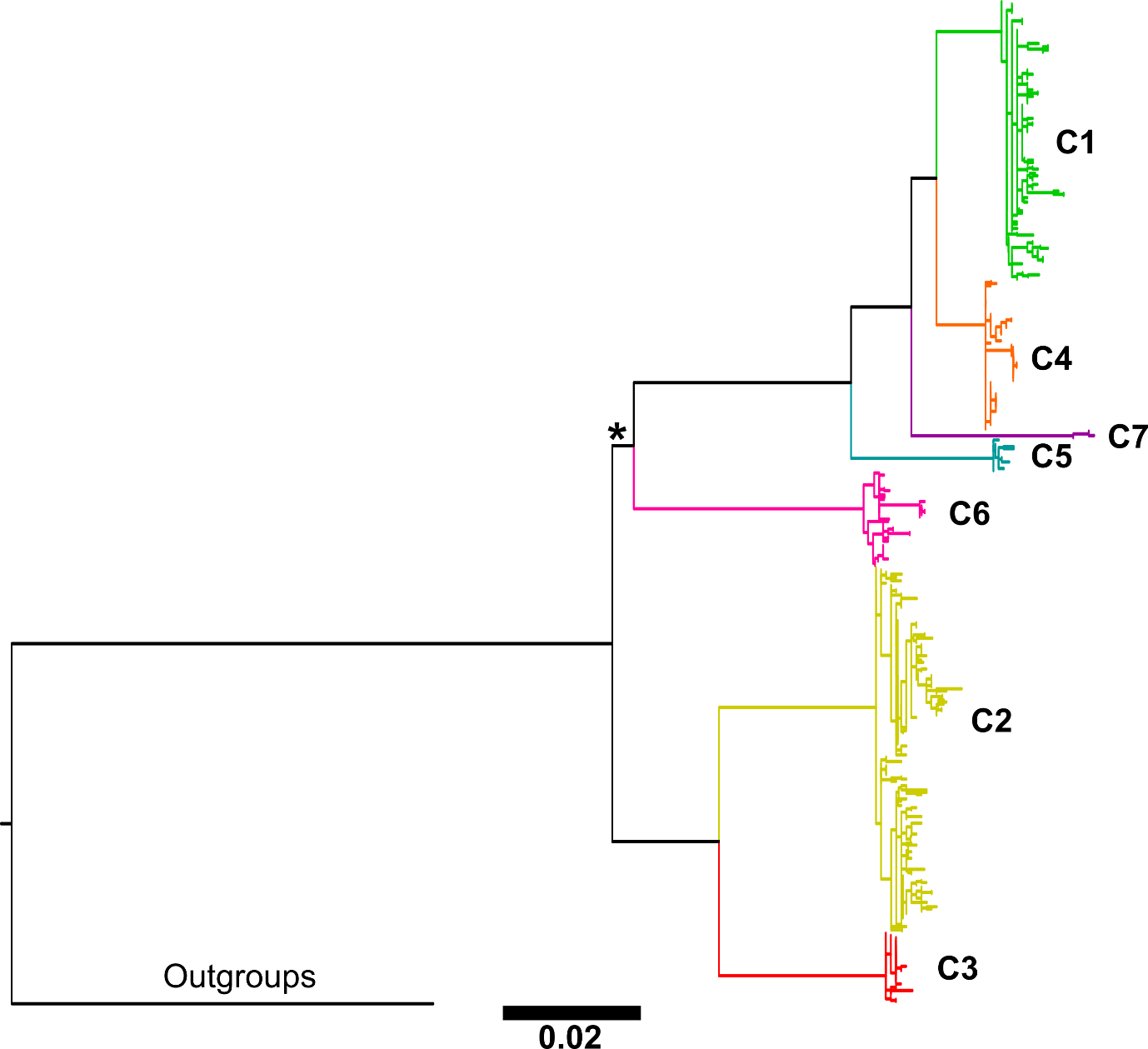

Figure S10. The combined (mtDNA+nDNA) based ML tree; “ * ” indicates weak support (BP < 70); all nodes without “ * ” were strongly supported (BP > 70).

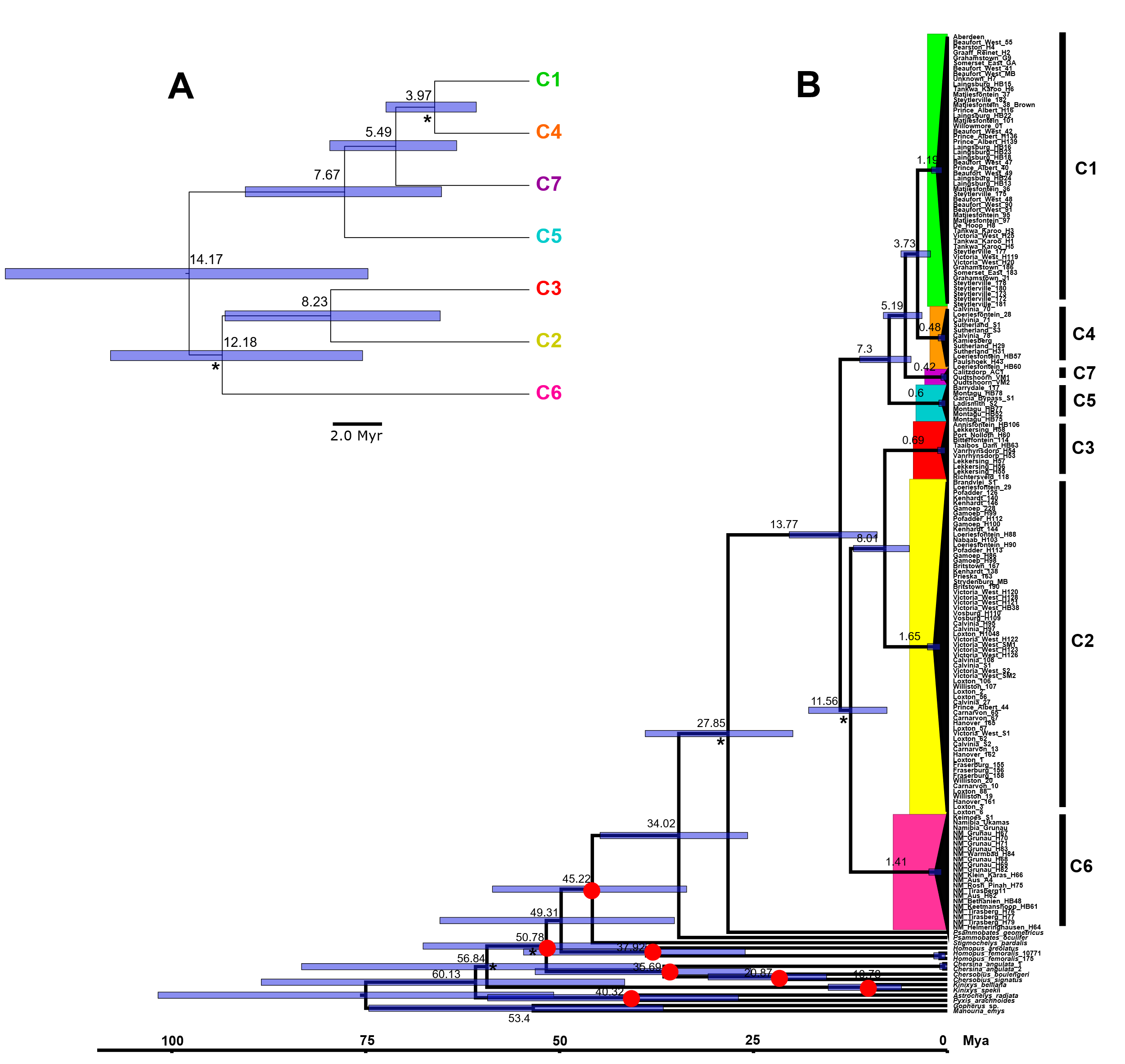

Figure S11. A: The time calibrated species tree inferred by Bayesian analyses with the StarBeast package in BEAST 2; B: The time calibrated gene tree generated by the StarBeast analysis; the seven red dots represent the seven constrained calibration nodes. On both the gene tree and species tree, the blue bar represents the estimated 95% HPD time range; the estimated age at each node is shown in the middle of each 95% HPD bar.

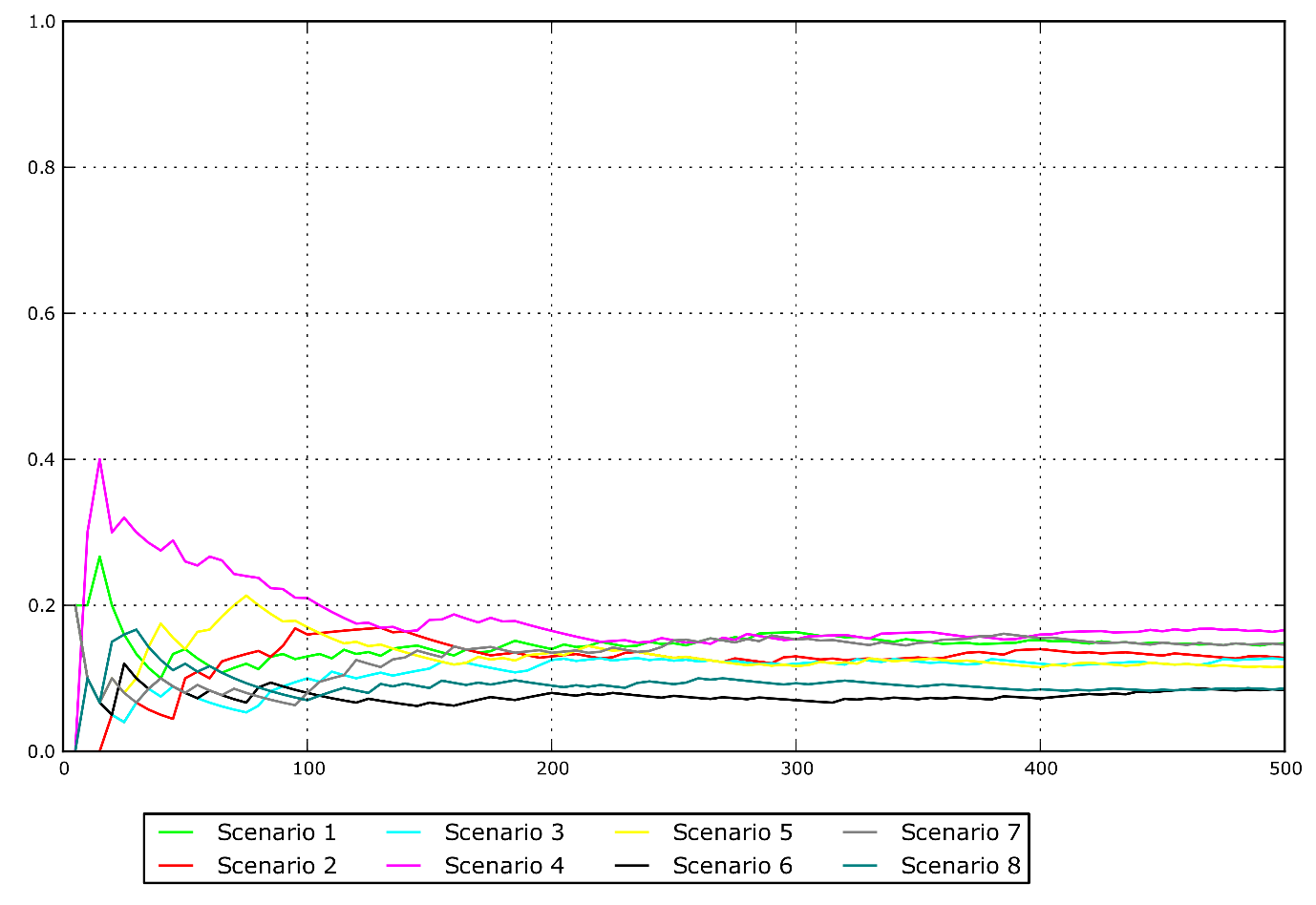

Figure S12. Best model selection using model comparison methods implemented in the ABC analyses. The scenario with highest Direct score (occupying the highest proportion) was considered the most plausible scenario.

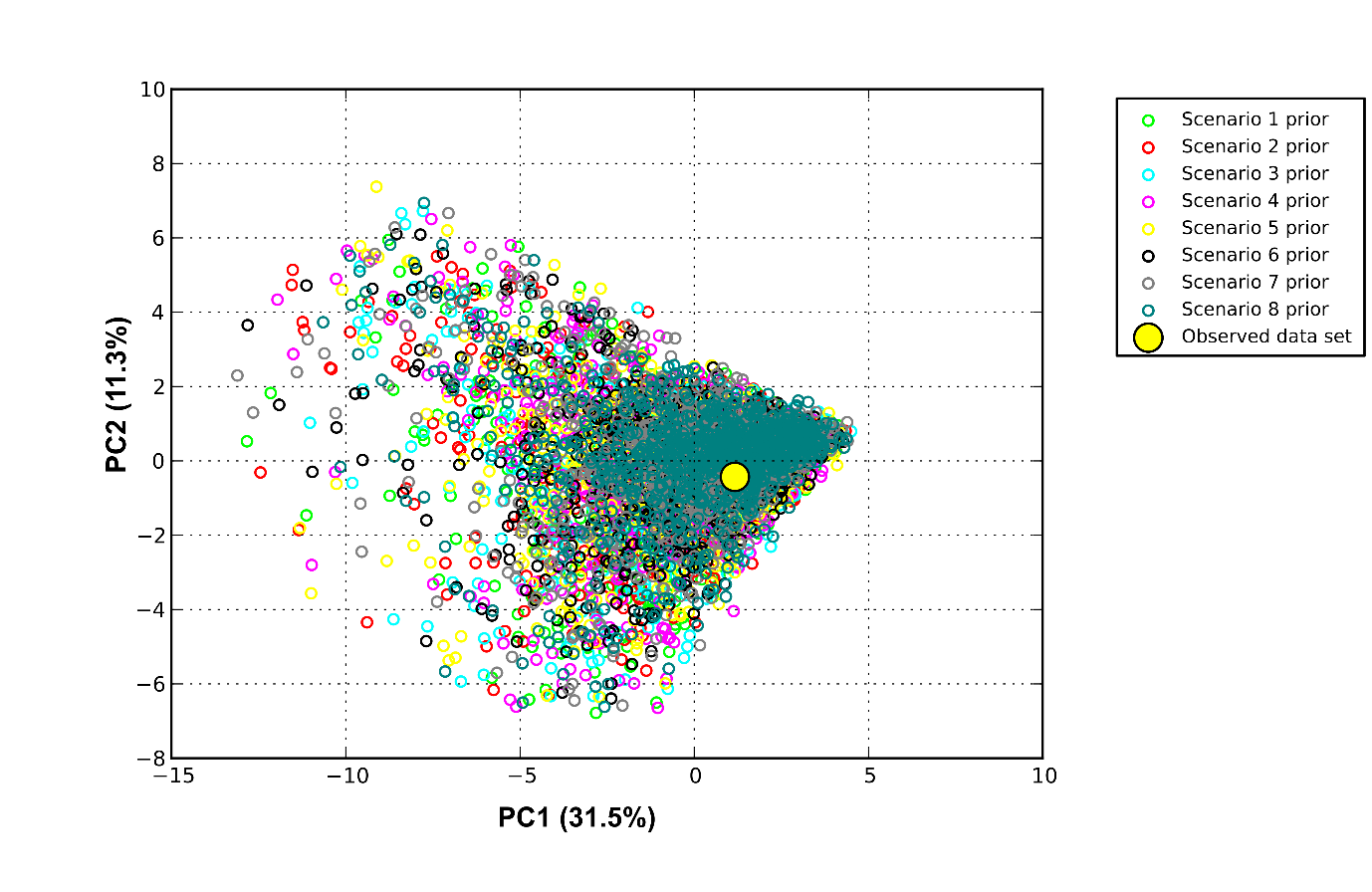

Figure S13. Scatterplot showing the PCA results evaluating the scenarios and priors in the ABC analyses. Each small dot represents a simulated dataset from the reference table and the large yellow dot represents the observed dataset. I expected the yellow dot to fall in the range of the small dots, which indicated that sampling was adequate.

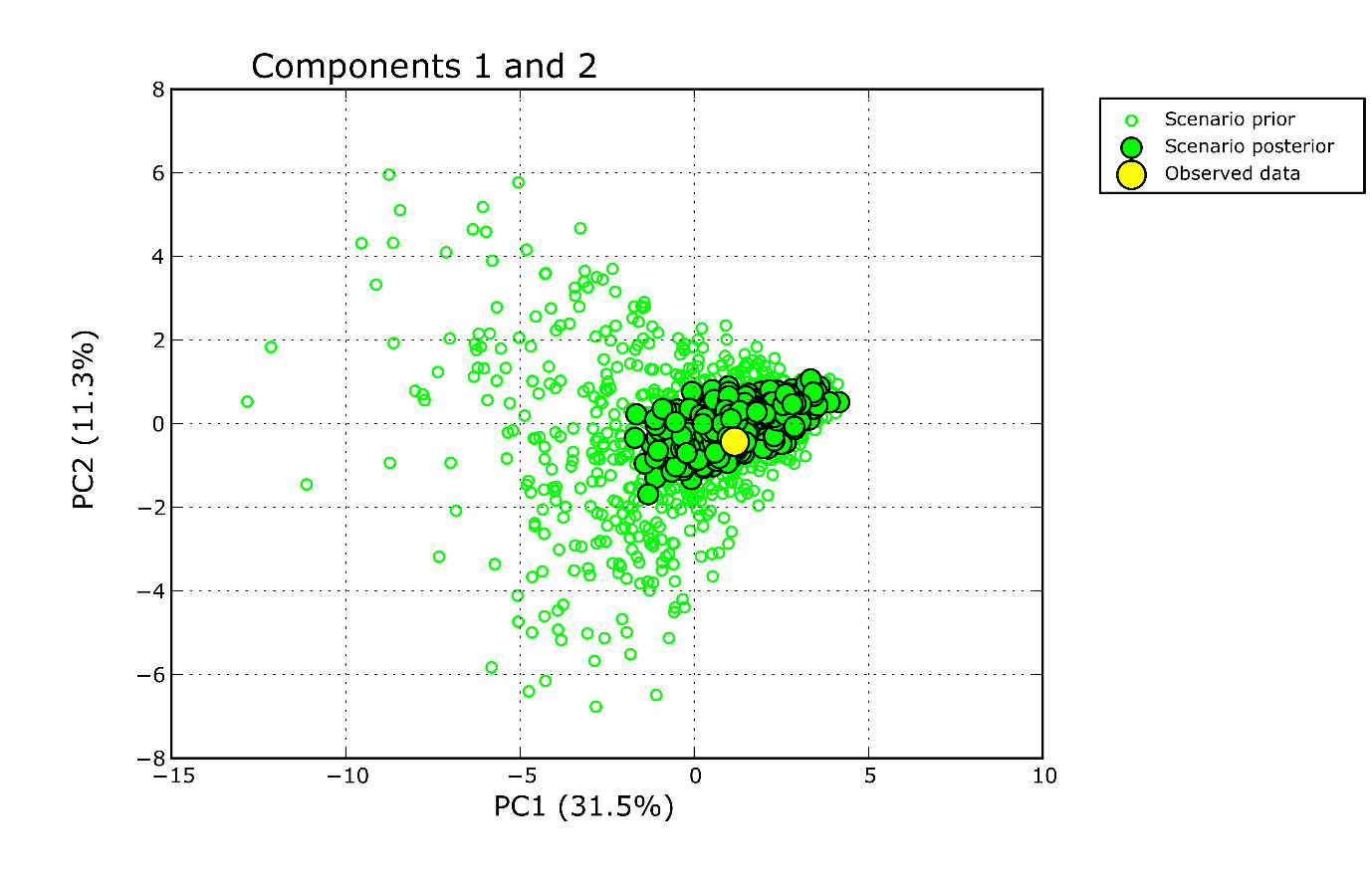

Figure S14. Results of the model checking ABC analyses. The observed dataset (shown as the big yellow dot) fell within the range of both the simulated priors (open dots) and posteriors (medium size dots) datasets, showing that sampling was adequate, and the simulation analysis results reliable.

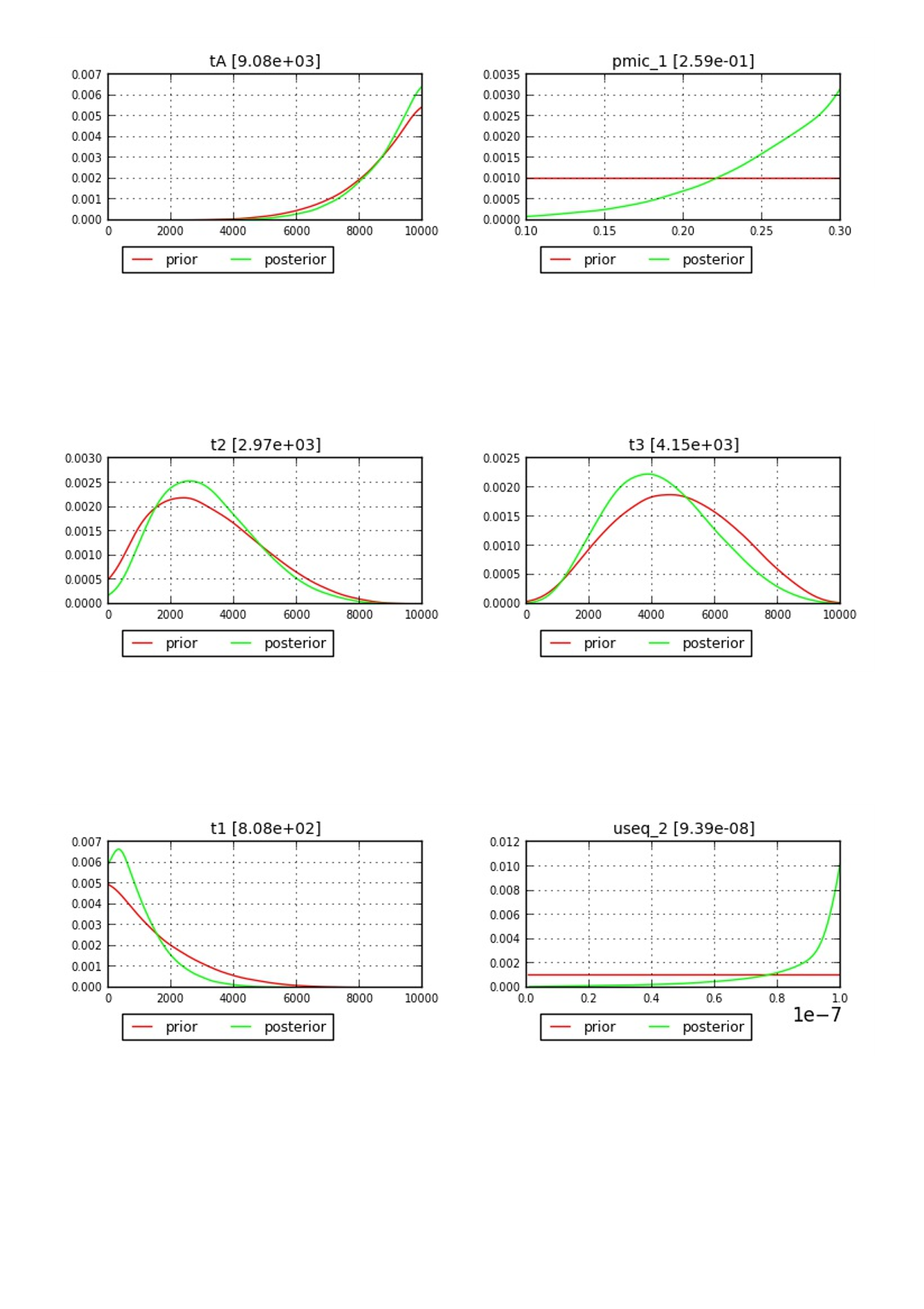

Figure S15. The parameters involved in the ABC analyses between priors and posteriors. The majority of the distributions of the priors and posteriors revealed a good match, implying that the simulation analysis results were reliable.

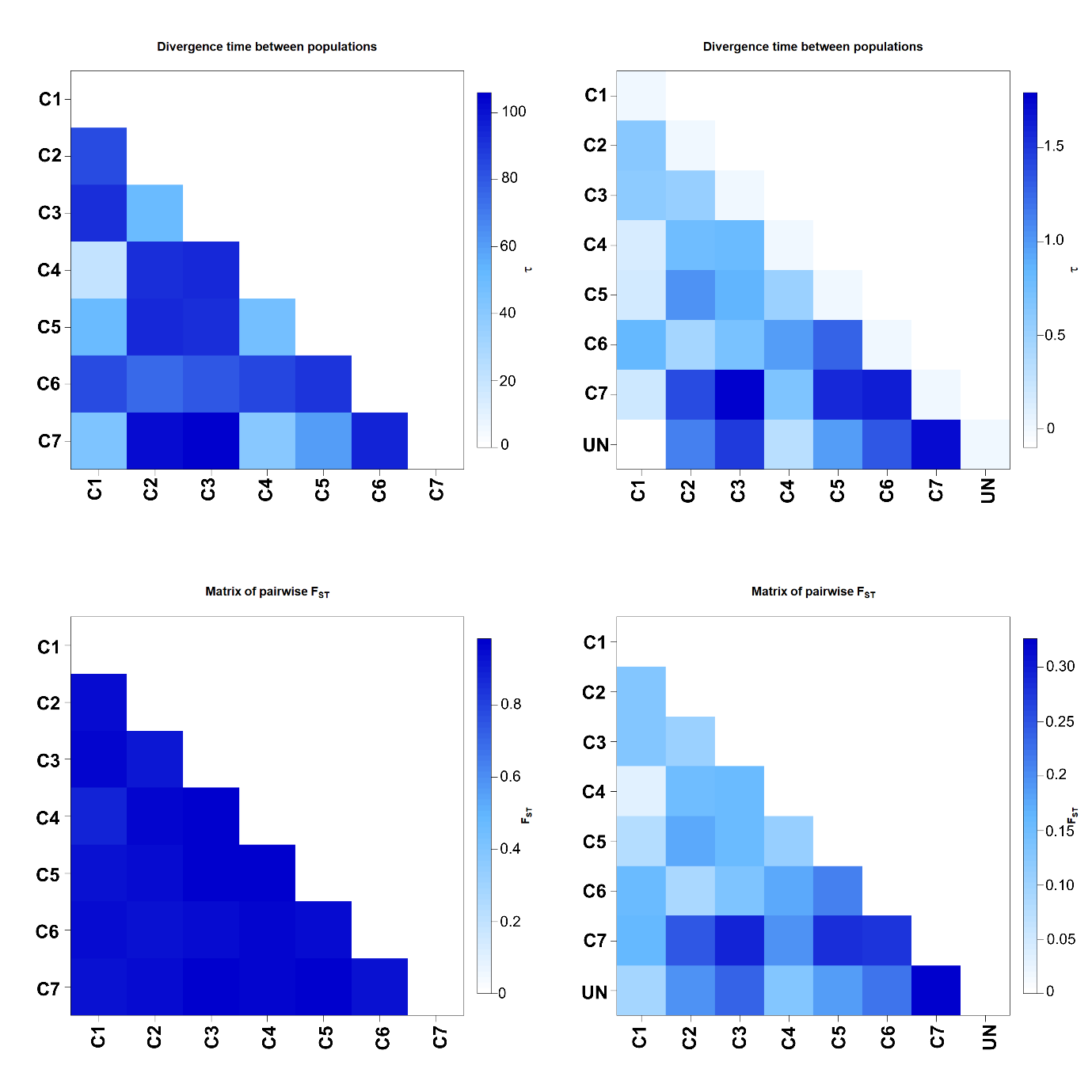

Figure S16. The relative divergence time (tau) matrix and the pairwise *F_ST_* matrix generated among clades from the AMOVA analyses between mtDNA (shown on the left side) and microsatellite DNA (shown on the right side) datasets. “UN” refers to the Uniondale population.
